# Analysis of genes within the schizophrenia-linked 22q11.2 deletion identifies interaction of *night owl/LZTR1* and *NF1* in GABAergic sleep control

**DOI:** 10.1101/755454

**Authors:** Gianna W. Maurer, Alina Malita, Stanislav Nagy, Takashi Koyama, Thomas M. Werge, Kenneth A. Halberg, Michael J. Texada, Kim Rewitz

## Abstract

The human 22q11.2 chromosomal deletion is one of the strongest identified genetic risk factors for schizophrenia. Although the deletion spans a number of genes, the contribution of each of these to the 22q11.2 deletion syndrome (DS) is not known. To investigate the effect of individual genes within this interval on the pathophysiology associated with the deletion, we analyzed their role in sleep, a behavior affected in virtually all psychiatric disorders, including the 22q11.2 DS. We identified the gene *LZTR1* (*night owl, nowl*) as a regulator of sleep in *Drosophila*. Neuronal loss of *nowl* causes short and fragmented sleep, especially during the night. In humans, *LZTR1* has been associated with Ras-dependent neurological diseases also caused by *Neurofibromin-1* (*Nf1*) deficiency. We show that *Nf1* loss leads to a night-time sleep phenotype nearly identical to that of *nowl* loss, and that *nowl* negatively regulates Ras and interacts with *Nf1* in sleep regulation. We also show that *nowl* is required for metabolic homeostasis, suggesting that *LZTR1* may contribute to the genetic susceptibility to obesity associated with the 22q11.2 DS. Furthermore, knockdown of *nowl* or *Nf1* in GABA-responsive sleep-promoting neurons elicits the sleep-fragmentation phenotype, and this defect can be rescued by increased GABAA receptor signaling, indicating that Nowl promotes sleep by decreasing the excitability of GABA-responsive wake-driving neurons. Our results suggest that *nowl*/*LZTR1* may be a conserved regulator of GABA signaling and sleep that contributes to the 22q11.2 DS.

## Introduction

Recent genome-wide association studies have identified chromosomal deletions and duplications that confer elevated risk of neuropsychiatric disorders [1, 2]. One of these, the 22q11.2 deletion, which spans 43 genes and occurs in 1 of every 2-4,000 births worldwide [3], is the most prominent known genetic risk factor for development of schizophrenia and is associated with high risk of neuropsychiatric disease [4, 5]. This deletion leads to a variety of phenotypes, including clinical manifestations such as congenital defects of the palate and heart, learning and cognitive disabilities, and sleep disturbances [6]. Sleep abnormalities are among the most common manifestations of neuropsychiatric disease [7], with up to 80% of autism and schizophrenia patients experiencing sleep disturbances [8, 9]. Individuals affected by the 22q11.2 deletion have a roughly 25-30% chance of developing schizophrenia or other psychiatric disorders that include sleep disturbances [10]. However, the possible contribution of each of the individual genes spanned by this deletion to this array of symptoms is not clear. Although certain disease phenotypes have been linked to single genes, such as *TBX1*, found to account for cardiac defects observed in 22q11.2 deletion carriers [11], the genes contributing to the 22q11.2-linked behavioral phenotypes such as sleep disturbance are not characterized.

Many schizophrenic patients exhibit abnormal sleep patterns such as delayed sleep onset, difficulty in maintaining sleep, and reduced overall amount of sleep [9]. Thus, genes that might play a role in schizophrenia and other behavioral symptoms of the 22q11.2 DS can potentially be identified by their sleep phenotypes. Sleep is an evolutionarily ancient conserved behavior that can be studied in lower organisms such as the fruit fly *Drosophila melanogaster*, a commonly used invertebrate model. *Drosophila* sleep has been rigorously examined and found to exhibit several of the mammalian hallmarks of sleep, such as sustained periods of quiescence, increased arousal threshold, and circadian and homeostatic regulation [12, 13]. Indeed, recent studies in *Drosophila* have identified genes associated with human neurodevelopmental and psychiatric disorders that disrupt sleep and circadian rhythm, such as the candidate autism-spectrum-disorder gene *Cullin-3* (*Cul3*) [14, 15] and the candidate gene *alicorn* (*alc*) linked to the 1q21.1 deletion, another CNV that confers schizophrenia risk [16].

To identify genes that contribute to the behavioral deficits of the 22q11.2 DS, we examined the role in sleep of individual genes within this interval by knockdown of their *Drosophil*a homologs in the nervous system of the fly. We identified *night owl* (*nowl, CG3711*), an ortholog of the human *LZTR1* (*leucine-zipper-like transcription regulator 1*), as a modulator of night-time sleep in *Drosophila*. In humans, mutations in *LZTR1* cause neurofibromatosis, a genetic disorder characterized by the growth of benign nervous-system tumors, that is most commonly known to result from mutations in *Neurofibromin-1* (*Nf1*) [17, 18]. Here, we show that neuronal knockdown or mutation of *nowl* causes sleep disturbances including highly increased fragmentation of sleep and reduced total night-time sleep, similar to neuronal loss of *Neurofibromin-1* (*Nf1*) in *Drosophila*. Nf1 is a negative regulator of the Ras signaling pathway and an activator of cAMP signaling [19, 20]; little is known about the function of *LZTR1*, except that its loss has been associated with neurofibromatosis. Interestingly, we find that *nowl* and *Nf1* inhibit Ras signaling and interact genetically in the regulation of sleep in *Drosophila*. We find that *nowl* acts through gamma-aminobutyric acid (GABA)-responsive neurons expressing the ionotropic GABA receptor Rdl to maintain sleep. Furthermore, the sleep fragmentation exhibited by *nowl* mutant flies can be rescued by increased inhibitory signaling through this receptor. Because *nowl* knockdown phenocopies the sleep disturbances observed with neuronal *Cul3* knockdown, we suggest that Nowl may interact with Cul3 and Nf1 in GABA-responsive neurons to regulate sleep. Thus, we have identified the 22q11.2-linked gene *nowl*, orthologous to human *LZTR1*, as a regulator of GABAergic signaling and night-time sleep in *Drosophila*. We suggest that *LZTR1* is a promising candidate gene that may contribute to the sleep disturbances and behavioral symptoms seen in patients carrying the 22q11.2 deletion.

## Results

### *Drosophila* screening identifies an ortholog of human *LZTR1* as a regulator of sleep

Sleep disturbance is a common manifestation of neuropsychiatric disorders in general and of the 22q11.2 DS. To determine which, if any, of the individual genes within this deletion might be required for normal sleep, we first identified the *Drosophila* orthologs of the human genes within the 22q11.2 interval. From the 43 genes lying within the 22q11.2 deletion, we identified 26 highly conserved orthologs in *Drosophila* (with a DIOPT score ≥ 6) [21] (Fig. 1A). Next, we conducted an RNA-interference (RNAi)-based screen to investigate whether the neuronal function of these *Drosophila* genes might be important for sleep, using the *Drosophila* Activity Monitor (DAM) system (TriKinetics), an automated beam-crossing locomotion assay that is the widely accepted standard for *Drosophila* sleep studies. In these assays, locomotor quiescence of longer than 5 minutes is taken to indicate a sleep-like state, comparable to mammalian sleep [12, 13]. To determine whether each identified *Drosophila* ortholog might play a role in sleep, we knocked each gene down using the nervous system-specific *elav-GAL4* (*elav>*) line to drive RNAi expression. When possible, two independent RNAi lines targeting each gene were tested. The effectiveness of RNAi knockdown was enhanced by the co-expression of *Dicer-2* (*Dcr-2*) [22]. Some RNAi constructs induced lethality when expressed pan-neuronally or displayed a non-inflating wing phenotype due to a gene disruption at the insertion site of the transgene [23]. These constructs were excluded from further analysis.

**Figure 1.**
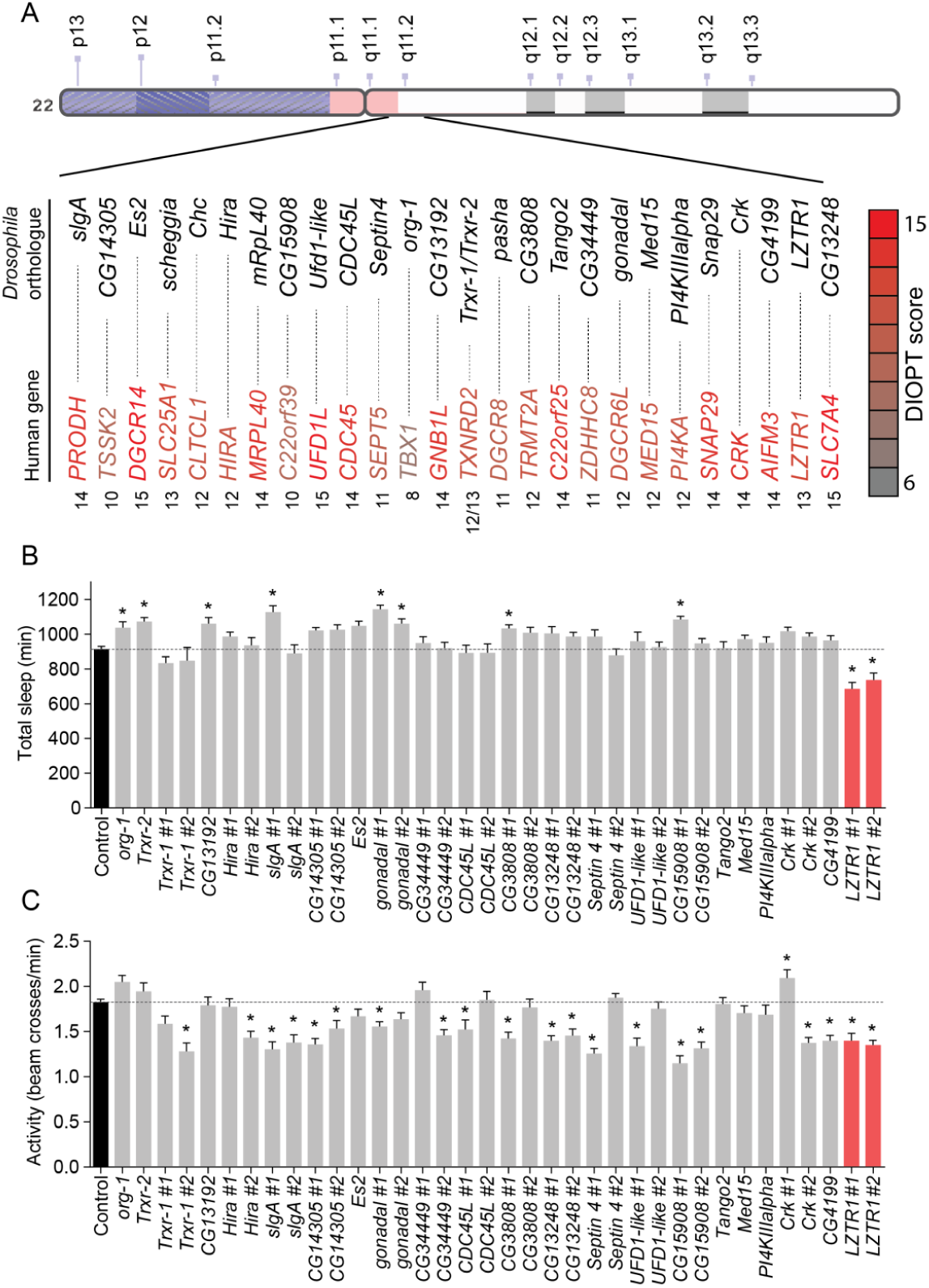
Screening *Drosophila* orthologs of human genes within the 22q11.2 deletion identifies *LZTR1* as a modulator of sleep and activity in *Drosophila*. (A) Schematic diagram of the human 22q11.2 deletion. For the 43 genes spanned by the 22q11.2 deletion, 26 orthologs were identified in *Drosophila melanogaster* with a DIOPT conservation score ≥ 6. (B, C) Total sleep and activity in adult males with pan-neuronal *RNAi*-mediated knockdown of each of these *Drosophila* orthologs. The *elav-GAL4* (*elav>*) pan-neuronal driver line was crossed to *UAS-RNAi* transgenes targeting individual gene orthologs, with n=16 flies for each genotype. For controls, *elav>* was crossed to *w*^*1118*^, the genetic background for the RNAi lines, with n=140 flies. When possible, two independent *UAS-RNAi* transgenes were used against each gene. RNAi efficiency was enhanced by co-expressing *UAS-Dicer-2* (*Dcr-2*). Total sleep (minutes) of 3-to-7-day-old males was recorded over a 24-hour period (B). Average activity was measured as the number of beam crosses per minute (C). Knockdown of *LZTR1* in the nervous system reduces both total sleep and locomotor activity. Graphs represent means with SEM. Statistical significance was determined using a one-way ANOVA with Dunnetts post-hoc testing (* p<0.05).

Pan-neuronal knockdown of several of the *Drosophila* 22q11.2 gene orthologs caused changes in total sleep and activity in adult males (Fig. 1B and 1C). Most of these manipulations led to a decrease in locomotor activity, with unchanged or increased sleep amount. Nervous system-specific knockdown of *CG3808* (human *TRMT2A*), *CG15908* (*C22orf39*), *gonadal* (*DGCR6L*), *org-1* (*TBX1*), or *slgA* (*PRODH*) increased the total amount of sleep exhibited by males, while only the knockdown of *CG3711*, the *Drosophila* ortholog of *LZTR1*, caused a decrease in total sleep (Fig. 1A). In contrast, in females, neuronal knockdown of several genes led to decreased sleep, including *CDC45L* (human *CDC45*), *CG13192* (*GNB1L*), *CG13248* (*SLC7A4*), *Crk* (*CRK*), *Es2* (*DGCR14*), *Septin4* (*SEPT5*), and *Ufd1-like* (*UFD1L*) (S1A Fig.). Locomotor activity was affected upon knockdown of several 22q11.2 genes, with most leading to hypoactive phenotypes in both males and females (Fig. 1C and S1B Fig.). We decided to focus our further studies on *LZTR1*, the only tested gene whose knockdown led to reduced total sleep in combination with decreased locomotor activity in males.

### *nowl* is required for night-time sleep

To investigate the role of *LZTR1* in sleep, we examined the daily sleep pattern in animals with neuronal knockdown of *LZTR1*. In light of the reduced-night-time-sleep phenotype displayed by these animals, we chose to refer to this gene as *night owl* (*nowl*) (Fig. 2A). We found that nervous-system-specific knockdown of *nowl* (*elav>nowl-RNAi*) reduced night-time sleep compared to controls (*elav>+* and *nowl-RNAi/+, i.e., elav>* and *nowl-RNAi* crossed to the *w*^*1118*^ genetic background for the RNAi line). To further investigate the effect of *nowl* loss, we examined the sleep phenotype of a homozygous-viable transposon-insertion mutant of *nowl, Mi{ET1}CG3711*^*MB12128*^ (for simplicity we hereafter refer to this allele as *nowl*^*1*^), which carries a 6-kb insertion into the third exon of *nowl* [24] (Fig. 2B-2D). Transcript levels of *nowl* have previously been shown to be drastically reduced in this mutant [25]. Like animals with neuronal knockdown, *nowl*^*1*^ mutants exhibited decreased sleep, primarily during the second part of the night, compared with the genetic-background controls (*w*^*1118*^*/Y* and *+/Y*, where “+” denotes a wild-type X chromosome and “Y” a wild-type Y chromosome). When sleep data was binned into day and night periods, it showed that *nowl* loss-of-function had a much larger effect on sleep during the night than during the day (Fig. 2E-2G). Males, but not females, exhibited a slight decrease in daytime sleep. On the other hand, night-time sleep dropped dramatically in both males and females. Loss of one copy of *nowl* (on the X chromosome) in females did not elicit a phenotype, suggesting that the *nowl*^*1*^ mutation is a recessive loss-of-function allele, consistent with the transcript-level defect previously reported [25]. To further confirm these results, we generated a CRISPR/Cas9-mediated deletion of the *nowl* coding sequence. Like neuronal knockdown and the *nowl*^*1*^ mutation, this deletion mutant (*nowl*^*KO*^) showed reduced night-time sleep (S2A Fig.). Taken together, these data indicate that *nowl* is important for proper sleep behavior, mainly during the night.

**Figure 2.**
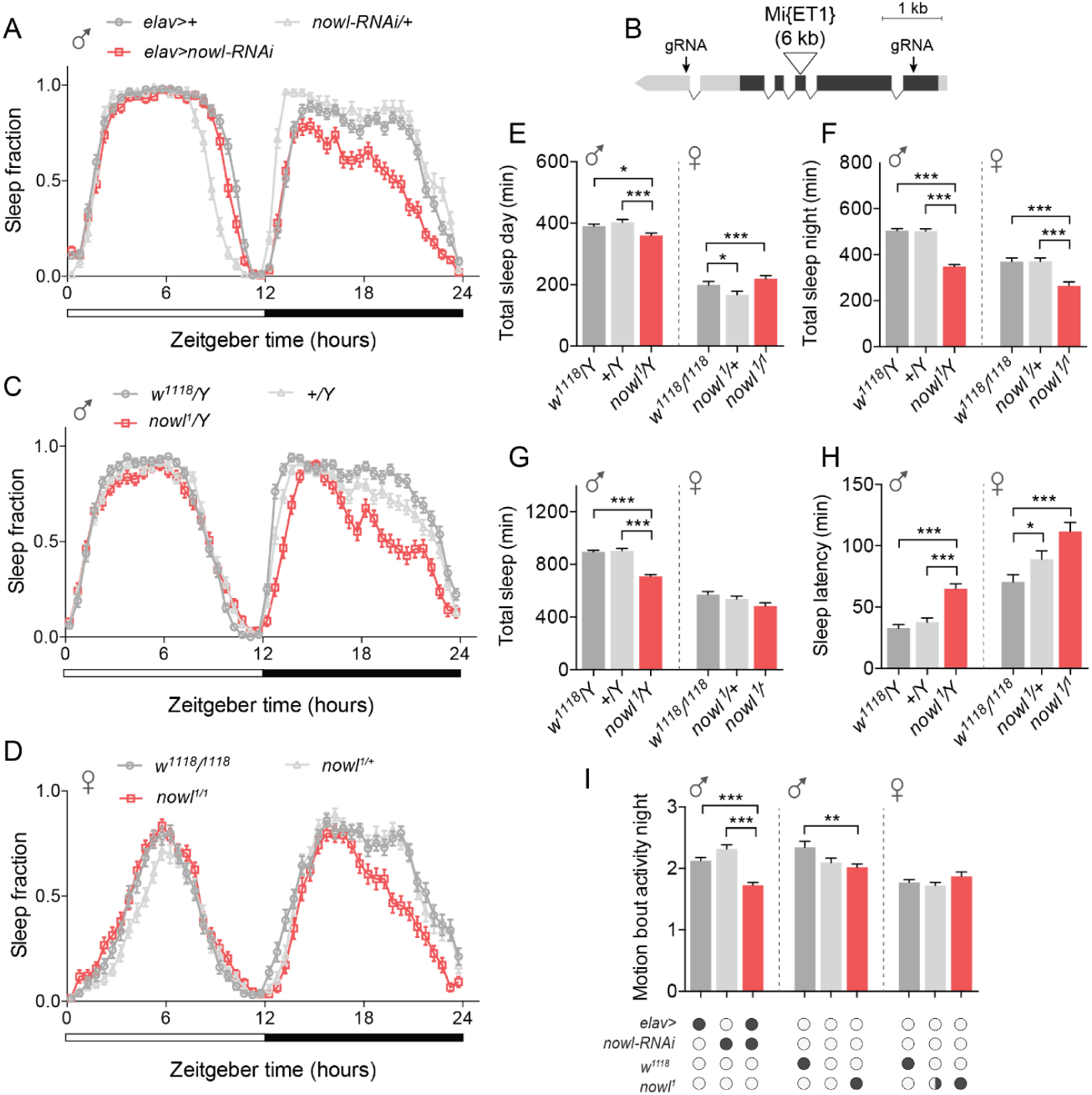
Neuronal loss of the *LZTR1* homolog *night owl* (*nowl)* reduces night-time sleep in *Drosophila*. (A) Daily sleep profiles across a 12-hour light, 12-hour dark (white and black bars) cycle for controls (*elav>%* and *nowl-RNAi/%*) compared to *elav>nowl-RNAi*. Sleep data are binned into 30-minute intervals. (B) Schematic of the *nowl* locus. The gene comprises 5′ and 3′ untranslated regions (light grey) and a multi-exon coding region (dark grey). The locations of the 6-kb *Mi{ET1} Minos* transposable element inserted in the *nowl*^*1*^ allele and the guide-RNA (gRNA) target sites used to induce the *nowl*^*KO*^ CRISPR/Cas9-mediated lesion are indicated. (C, D) Sleep profiles for (C) male *nowl*^*1*^*/Y* and (D) female *nowl*^*1*^*/nowl*^*1*^ mutant flies, compared to control genotypes (in C: *w*^*1118*^ *nowl*^*%*^*/Y* and *w*^*%*^ *nowl*^*%*^*/Y*; in D: *w*^*1118/1118*^ and *nowl*^*1*^*/%*). (E-F) Total day-and night-time sleep (minutes) in male and female flies. (E) Total daytime sleep is decreased in male *nowl*^*1*^ mutant flies, but not in female homozygous mutant flies. (F) Total nighttime sleep is significantly decreased in hemizygous male and homozygous female *nowl*^*1*^ mutants, but not in heterozygous females. (G) Overall total sleep (min.) in male and female *nowl*^*1*^ mutant flies compared to controls. Loss of *nowl* in male flies significantly decreases total sleep compared to controls. (H) Sleep latency measured for male and female *nowl*^*1*^ mutant flies compared to controls. Loss of *nowl* in both male and female flies significantly increases sleep latency compared to controls. (I) Quantification of motion-bout activity (min.) during night. Motion-bout activity during the night is significantly reduced in *nowl*-knockdown flies. Graphs represent means with SEM (n=32-92). For normally distributed data, significance was determined using a one-way ANOVA with Dunnetts post-hoc testing. For non-normally distributed data, Kruskal-Wallis test with Dunns post-hoc testing was used (* p<0.05, ** p<0.01, *** p<0.001).

Although we observed a decrease in sleep levels overall, sleep levels in these flies still dropped in anticipation of light-dark and dark-light transitions, consistent with an intact circadian clock. To investigate whether *nowl* mutant males do maintain a normal circadian rhythm, adult animals entrained to a 12/12-hour light-dark cycle were kept under constant darkness (free-running conditions) for 8 days. The average circadian period of *nowl*^*1*^ mutants during these days was 23.5 hours, which is very similar to the corresponding controls’ free-running periods of 23.3 and 23.4 hours (S3A and S3B Fig.). This indicates that the *nowl* mutants can entrain to and maintain a normal circadian rhythm, suggesting that *nowl* function is not required for proper functioning of the circadian clock.

### Loss of *nowl* disrupts sleep architecture and phenocopies lack of *Cul3* and *Nf1*

Sleep-onset difficulties are common in psychiatric disorders [26]. Interestingly, both male and female *nowl* mutants exhibit increased sleep onset-latency, defined as the time between lights-off and the first sleep bout; that is, they take longer to fall asleep than controls (Fig. 2H and S2B Fig.). Control males’ latency was 40 minutes, while hemizygous *nowl* mutants on average took 70 minutes to fall asleep after lights-off. Homozygous *nowl-*mutant females also exhibited delayed sleep onset and on average remained active for 110 minutes before their first sleep episode, while control females fell asleep after 70 minutes. Sleep latency was also slightly increased in heterozygous *nowl* mutant females, in which the time to sleep onset was 90 minutes, intermediate between controls and homozygotes.

To confirm that the observed phenotype was related to sleep *per se* and not to altered activity patterns, we analyzed average activity during periods in which the animals were active in the night. Motion-bout activity was significantly reduced when *nowl* was knocked down in the nervous system compared to driver- and *UAS*-alone control genotypes (Fig. 2I), while *nowl* mutant flies also displayed reduced or unchanged motion-bout activity levels compared to control flies. This confirms that the observed sleep phenotype in animals lacking *nowl* is not the result of hyperactivity but rather is a specific sleep-disruption phenotype.

We next examined sleep fragmentation, which is associated with many psychiatric disorders and characterized by multiple short periods of sleep, *i.e.*, an increased number of shorter sleep bouts. In the initial screen of genes within the 22q11.2 deletion, we found that knockdown of some genes including *slgA (PRODH)* and *nowl/LZTR1* also altered the number and duration of sleep bouts (S4A-S4D Fig.). To further investigate whether loss of *nowl* plays a role in sleep maintenance following sleep initiation, we analyzed average sleep-bout number and length during day and night time in males lacking *nowl*. Knockdown of *nowl* in the nervous system using either of the two pan-neuronal drivers *elav>* and *nSyb*-*GAL4* (*nSyb>*) caused a strong increase in fragmentation of night-time sleep, with a large decrease in sleep-bout duration combined with an increase in the number of sleep bouts, that was not observed during daytime (Fig. 3A-3D). Mutant flies carrying the *nowl*^*1*^ loss-of-function allele or the CRISPR-induced *nowl*^*KO*^ deletion exhibited similar sleep-fragmentation patterns (S2C and S2D Fig.).

**Figure 3.**
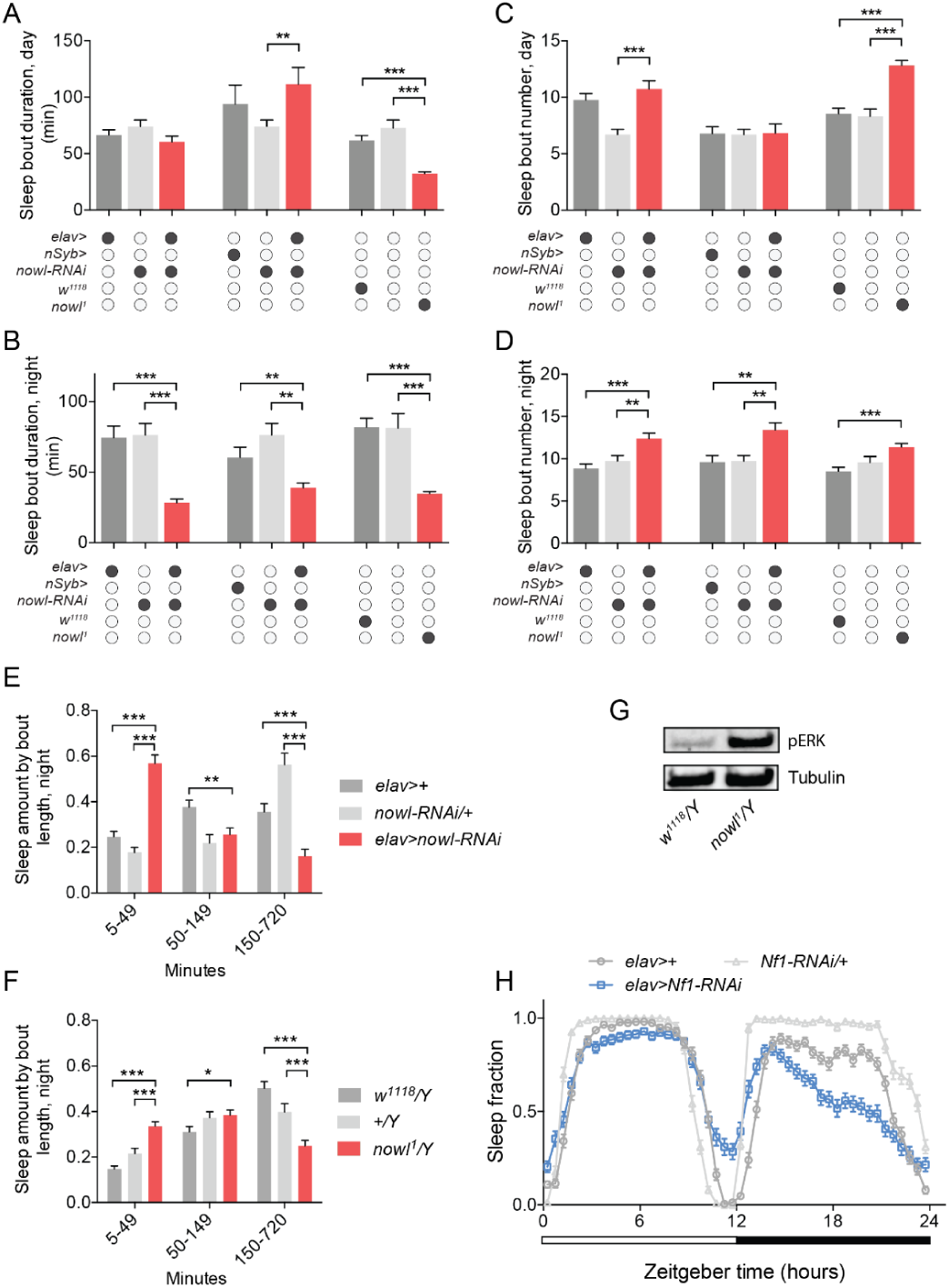
Loss of *nowl* in the nervous system causes sleep fragmentation. (A-D) Quantification of average sleep-bout duration (A and B) and number (C and D) during day and night for males with pan-neuronal knockdown of *nowl* using the stronger (*elav>*) and weaker (*nSyb>*) GAL4 drivers and the *nowl*^*1*^ insertional mutant. Neuronal knockdown of *nowl* using *elav>* (*elav>nowl-RNA*) and *nSyb>* (*nSyb>nowl-RNAi*) or mutation disrupting *nowl* (*nowl*^*1*^*/Y*) result in decreased sleep-bout length and increased sleep-bout number, with a stronger effect during night-time, compared to controls (*elav>+, nowl-RNAi/+, nSyb>/+, w*^*1118*^*/Y*, and *+/Y*). (E-F) Fraction of time male animals spent sleeping in short (5-49-min.), medium (50-149-min.) or long (150-720-min.) sleep bouts during night for controls (*elav>+* and *nowl-RNAi/+*) compared to *elav>nowl-RNAi* animals and for *nowl*^*1*^ (*nowl*^*1*^*/Y*) mutants compared to controls (*w*^*1118*^*/Y* and *+/Y*). Flies carrying a *nowl* mutation or in which *nowl* has been knocked down neuronally spend significantly less time in long consolidated sleep episodes, compared to controls. (G) Western-blot analysis shows increased levels of phospho-ERK (pERK) in *nowl* mutant males compared to controls (*w*^*1118*^*/Y*), standardized to alpha-Tubulin. (H) Sleep pattern of *Neurofibromin-1* (*Nf1*) knockdown males under a 12-hour light/dark (white and black bars) cycle in 30-minute intervals shows effects on sleep compared to controls (*elav>+* and *Nf1-RNAi/+*) similar to animals lacking *nowl*, with decreased sleep mainly during the night. Graphs represent means with SEM (n=32-92). For normally distributed data, significance was determined using a one-way ANOVA with Dunnetts post-hoc testing. For non-normally distributed data, Kruskal-Wallis test with Dunns post-hoc testing was used (* p<0.05, ** p<0.01, *** p<0.001).

To more closely examine the sleep changes associated with night-time sleep fragmentation, we analyzed the distribution of sleep episodes by binning them by duration and calculating the time spent in each bin, for flies with reduced *nowl* expression in the nervous system and for *nowl* mutants, compared to control genotypes. Neuronal knockdown of *nowl* significantly increased the amount of time spent in short sleep bouts of 5-49 minutes, compared to control flies, which consolidated most of their night-time sleep into longer bouts of 150-720 minutes (Fig. 3E). A similar change was observed in *nowl* mutant flies (Fig. 3F). These data indicate that *nowl* is required for maintenance of proper sleep architecture in *Drosophila*.

Human *LZTR1* was recently shown to function through the Cul3-based ubiquitin ligase complex, as a component of which it is suggested to mediate substrate specificity [27, 28]. Interestingly, *Cul3* has also been implicated as a causative gene in psychiatric disorders [29] and has been shown to play an important role in *Drosophila* sleep [14, 15, 30]. We wondered whether *nowl*, like its ortholog *LZTR1*, might exert its effects on sleep through an interaction with *Cul3*. We therefore knocked down *Cul3* pan-neuronally and analyzed sleep patterns, and found that *Cul3* silencing in the nervous system caused a significant decrease in average sleep-bout duration and a significant increase in average sleep-bout number (S5A-S5D Fig.), during both day and night. This phenotype is similar to the sleep disruption caused by neuronal knockdown of *nowl*, which is consistent with an interaction between the two genes.

Mutations in human *LZTR1* have been linked with schwannomatosis, a form of neurofibromatosis, a neurological disorder associated with sleep problems, mental disabilities, and psychiatric disorders that can also be caused by mutation of the *Nf1* gene [17]. *Nf1* encodes a GTPase that negatively regulates the Ras signaling pathway. We therefore asked whether *nowl*, like *Nf1*, affects Ras signaling, by measuring the levels of phosphorylated ERK (pERK), a downstream effector of Ras and marker of Ras-pathway activation. We found increased levels of pERK in heads from *nowl* mutants (Fig. 3G), indicating that the Ras pathway is overactivated. Thus, the Nowl protein does appear to be required for normal Ras-pathway inhibition, either directly, like Nf1, as a negative regulator of Ras, or indirectly. We next investigated whether *Nf1* regulates sleep and found that neuronal knockdown of *Nf1* reduces night-time sleep nearly identical to loss of *nowl* (Fig. 3H). These similarities are consistent with a sleep-regulating genetic interaction between *nowl* and *Nf1*.

### Interactions between *nowl* and *Nf1* suggest that they act on a common sleep-regulatory pathway

To investigate the possibility of genetic interaction between *nowl* and *Nf1* in regulation of sleep, we overexpressed *Nf1* in *nowl-RNAi* animals. In this test, a rescue of the *nowl* short-sleep phenotype would indicate that *Nf1* functions downstream of *nowl* in a sleep-regulatory pathway. Neuronal overexpression of *Nf1* partially rescued the loss of night-time sleep and sleep fragmentation caused by *nowl* knockdown (Fig. 4A-4C), suggesting that *nowl* and *Nf1* may function in the same sleep-regulatory pathway, with Nf1 working downstream of Nowl. On the other hand, the double knockdown of *Nf1* and *nowl* produced a reduction of day-time sleep that was not observed by single RNAi directed against either *nowl* or *Nf1* alone (Fig. 4D). Next, we analyzed interaction by comparing effects in the single-hit knockdown compared to the double-hit knockdown of *nowl* and *Nf1*. As an initial step, we determined the efficiency and specificity of the RNAi-mediated knockdown using qPCR, and validated that the RNAi lines against *nowl* and *Nf1* both significantly reduced target gene expression (S6A and S6B Fig.). In animals with double knockdown of *nowl* and *Nf1* in the nervous system, the total daytime sleep amount decreased and fragmentation increased, compared to animals with single knockdown of either *nowl* or *Nf1* and to the control, indicating that *nowl* and *Nf1* interact in the regulation of daytime sleep (Fig. 4B). In animals with double knockdown of *nowl* and *Nf1*, total night-time sleep was reduced, similar to animals with single knockdown of *nowl* or *Nf1*, compared to the control (Fig. 4C). Simultaneous knockdown of *nowl* and *Nf1* also increased night-time sleep fragmentation compared to the control. However, the effect of the double knockdown on night-time sleep parameters largely resembled the effect of *Nf1* knockdown alone, suggesting an interaction between *nowl* and *Nf1* in night-time sleep regulation that may be epistatic. The ability of *Nf1* overexpression to partially rescue the loss of total night-time sleep and reduced sleep-bout duration during the night supports the notion that these two genes are functionally related and may converge on a common sleep-regulatory pathway.

**Figure 4.**
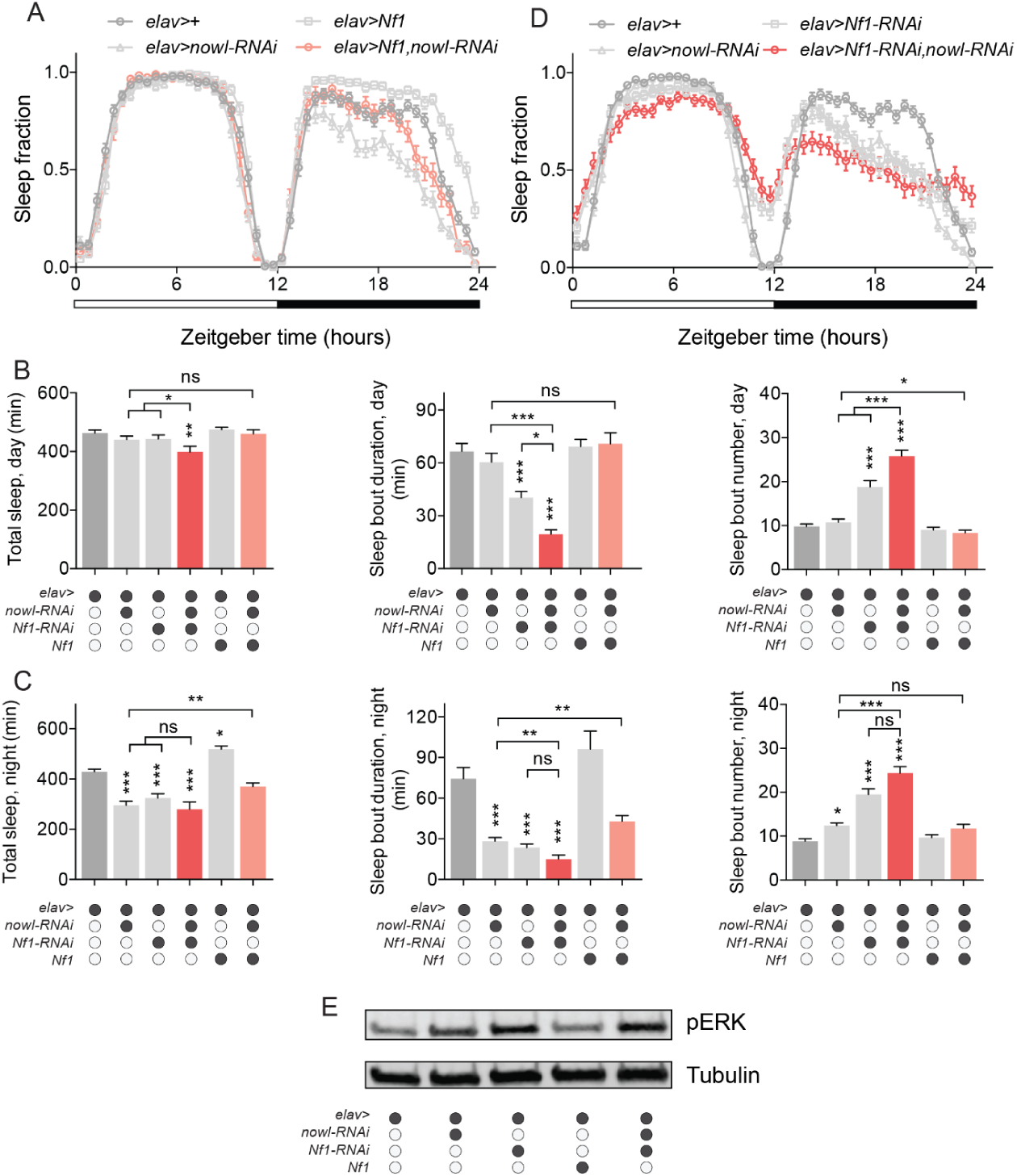
*nowl* and *Nf1* interact in the regulation of sleep. (A) Daily sleep profiles of animals under a 12-hour light/dark (white and black bars) cycle in 30-minute intervals. In *elav>Nf1, nowl-RNAi* animals (with neuronal knockdown of *nowl* and overexpression of *Nf1*), night-time sleep is partially restored compared to *nowl* knockdown alone. (B and C) Quantification of daytime (B) and nighttime (C) total sleep, sleep-bout duration, and sleep-bout numbers. The double knockdown (*elav>nowl-RNAi, Nf1-RNAi*) shows significant interactions between *nowl* and *Nf1* on day-time total sleep and sleep-bout duration and number, indicating an additive effect in the daytime. The effects of double knockdown on total nighttime sleep and nighttime sleep-bout duration and number are not different from *Nf1-RNAi* alone, suggesting an epistatic relationship between *nowl* and *Nf1* in night-time sleep regulation. *Nf1* overexpression partially rescues the *nowl-*knockdown effect on sleep. Controls are *elav>+*. (D) Daily sleep profiles of double-knockdown animals (*elav>Nf1-RNAi, nowl-RNAi*) display a pronounced decrease in sleep. Controls are *elav>+*. (E) Western blotting against phosphorylated ERK (pERK) shows that single and double knockdowns of *nowl* and *Nf1* cause increased pERK levels, standardized to alpha-Tubulin (controls are *elav>+*). Graphs represent means with SEM (n=32). For normally distributed data, significance was determined using a one-way ANOVA with Dunnett’s post-hoc testing. For non-normally distributed data, Kruskal-Wallis test with Dunn’s post-hoc testing was used (*p < 0.05, ** p < 0.01, *** p < 0.001).

Since our results indicated that Nowl inhibits the Ras pathway (Fig. 3G), we investigated the combined effect of *nowl* and *Nf1* loss on the activation of the Ras pathway. Our data show that silencing *nowl* or *Nf1* individually in the nervous system results in Ras overactivation, reflected in increased pERK levels (Fig. 4E). Consistent with this and with their genetic interaction in sleep regulation, the simultaneous knockdown of *nowl* and *Nf1* activates the Ras pathway to an extent similar to knockdown of *Nf1* alone. Together these effects indicate that *nowl* and *Nf1* interact to suppress Ras-pathway activity.

To investigate the genetic interaction between *nowl* and *Nf1* further, we compared the phenotypes of *Nf1* and *nowl* trans-heterozygous mutants with those of the individual heterozygous mutants. The *nowl* gene is on the X chromosome, and males are therefore hemizygous; heterozygous effects can only be studied in females. Female animals heterozygous for either *nowl* or *Nf1* displayed similar sleep patterns, with no significant loss of night-time sleep compared to the control (Fig. 5A-5C), whereas trans-heterozygous *nowl*^*+/-*^ *Nf1*^*+/-*^ mutant females exhibited a strong reduction in total sleep. Furthermore, trans-heterozygous *nowl*^*+/-*^ *Nf1*^*+/-*^ mutants displayed shorter sleep bouts, while animals heterozygous for mutations in either *nowl* or *Nf1* alone displayed no change in sleep-bout duration compared to controls (Fig. 5D-5G). This indicates that trans-heterozygous loss of both *nowl* and *Nf1* causes sleep fragmentation, further supporting the existence of a genetic interaction between *nowl* and *Nf1* and a functional relationship between these two genes in the regulation of sleep.

**Figure 5.**
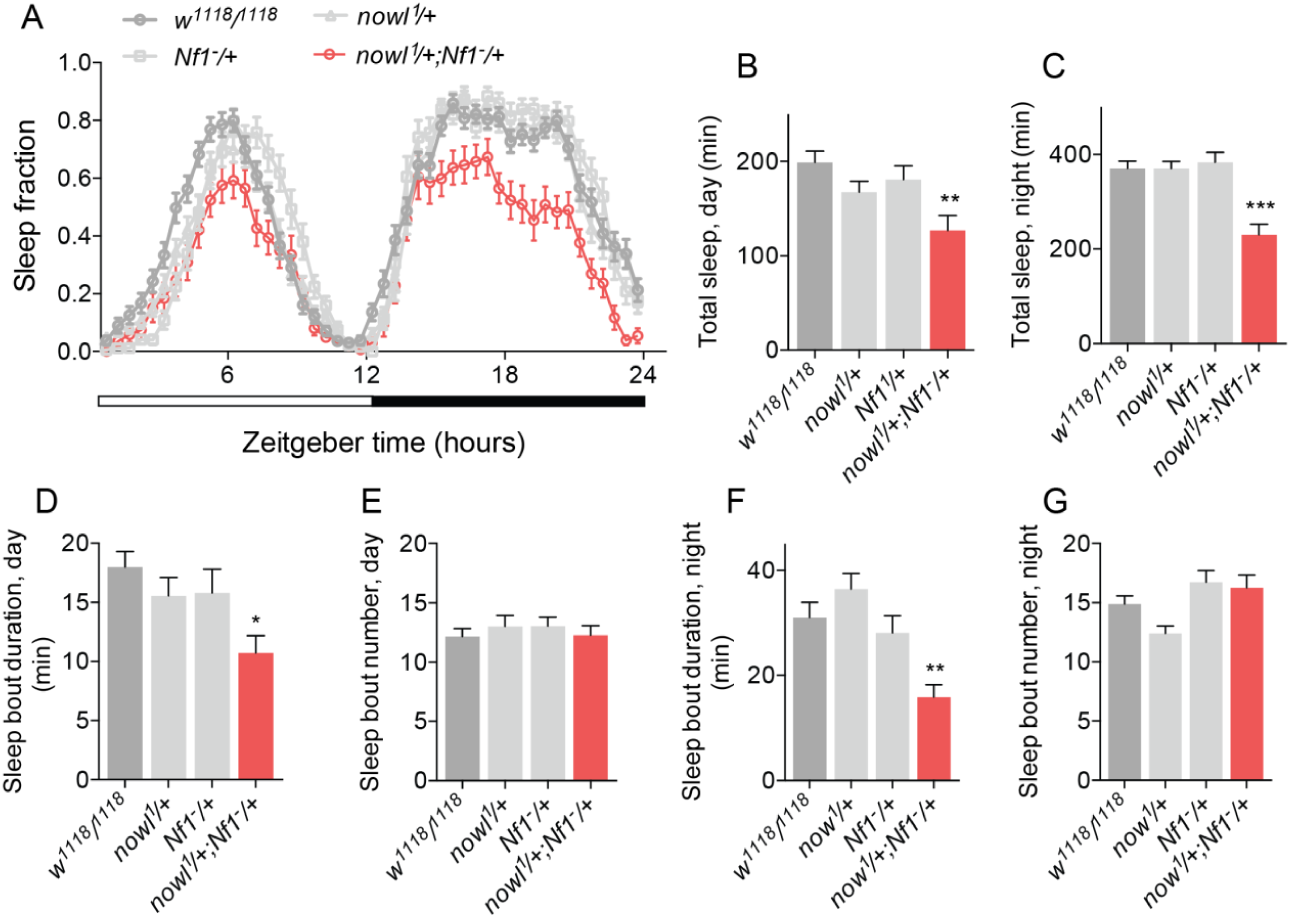
Interactions between *nowl* and *Nf1* suggest that they act on a common sleep-regulatory mechanism. (A) Sleep patterns of heterozygous *nowl*^*1*^*/+* and *Nf1*^*-*^*/+* females show that one copy of each is sufficient for maintaining proper sleep, while the trans-heterozygous (*nowl*^*1*^*/+*; *Nf1*^*-*^*/+*) mutants exhibit decreased sleep during both day and night. (B-G) Quantification of sleep parameters shows reduced average sleep-bout duration of trans-heterozygous *nowl*^*1*^*/+; Nf1*^*-*^*/+* females, indicating that *nowl* and *Nf1* interact to maintain sleep. Graphs represent means with SEM (n=32). For normally distributed data, significance was determined using a one-way ANOVA with Dunnett’s post-hoc testing. For non-normally distributed data, Kruskal-Wallis test with Dunn’s post-hoc testing was used (*p < 0.05, ** p < 0.01, *** p < 0.001).

### Neuronal *Nf1* and *nowl* are required to maintain metabolic homeostasis

Emerging evidence suggests that sleep is important for energy homeostasis and that insufficient sleep can lead to obesity [31]. A main function of sleep has been proposed to be an opportunity to replenish brain supplies of glycogen, which are depleted during periods of wakefulness [32, 33]. Since *nowl* and *Nf1* loss decrease night-time sleep and thereby increase wakefulness, we analyzed glycogen levels in these animals. Consistent with the notion that sleep is important for maintaining glycogen levels, we found that indeed glycogen levels were reduced in animals with neuronal knockdown of *nowl* and *Nf1* and in *nowl* mutants (Fig. 6A and 6B), which exhibits reduced night-time sleep. These data further suggest that neuronal function of *nowl* and *Nf1* is essential to maintenance of glycogen stores, which may be related to their importance for getting enough sleep. We next analyzed body triglyceride levels since the 22q11.2 deletion is associated with increased risk of obesity, which is also linked to insomnia [34, 35]. We found increased organismal levels of triglycerides in flies with neuronal knockdown of *nowl* and in *nowl* mutants (Fig. 6C), suggesting that they exhibit increased adiposity like human 22q11.2 carriers. Taken together our results suggest that *nowl* is required in the nervous system to maintain organismal energy homeostasis, a function that might be related to its role in sleep considering the vital role of sleep in regulation of energy homeostasis.

**Figure 6.**
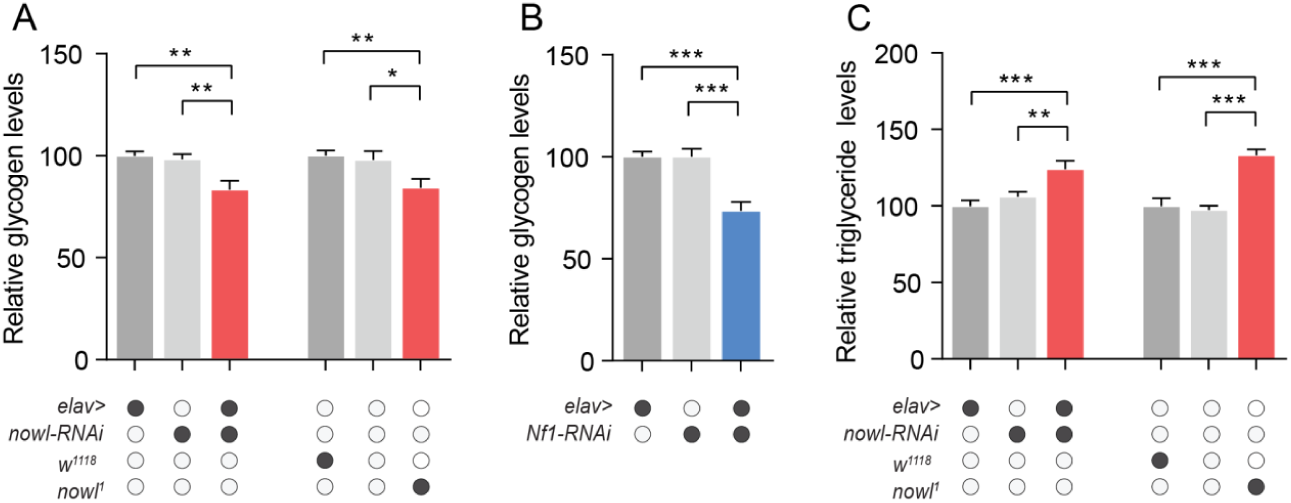
Loss *nowl* and *Nf1* leads to physiological changes that affect energy homeostasis. (A-C) Quantification of whole body glycogen (A,B) and whole body triglyceride (C) content in adult male flies. Neuronal knockdown of *nowl* (*elav>nowl-RNAi*) and *Nf1* (*elav>Nf1-RNAi*) or mutation disrupting *nowl* (*nowl*^*1*^*/Y*) results is reduced whole body glycogen levels (A, B) compared to controls (*elav>+, nowl-RNAi/+, Nf1-RNAi/+, w*^*1118*^*/Y*, and *+/Y*). (C) Whole body lipid content is increased animals with neuronal knockdown of *nowl* or a mutation disrupting *nowl*. Graphs represent means with SEM (n=12-20). For normally distributed data, significance was determined using a one-way ANOVA with Dunnett’s post-hoc testing. For non-normally distributed data, Kruskal-Wallis test with Dunn’s post-hoc testing was used (*p < 0.05, ** p < 0.01, *** p < 0.001).

### *Nf1* and *nowl* are required in GABA-responsive cells for night-time sleep

Changes in neuronal circuitry and neurotransmitter systems have been proposed to underlie several neuropsychiatric disorders. Signaling through GABA, in particular, has been found to be an important contributor to sleep in mammals [36]. GABAergic signaling has likewise been found to play a role in *Drosophila* night-time sleep initiation and maintenance [37-39]. Because we observed a strong disruption of both of these parameters in *nowl* knockdown and mutant flies, we wondered whether this gene might have a function in GABAergic signaling. To investigate this, we knocked down *nowl* in GABA-producing or GABA-responsive neurons using a panel of *GAL4* driver lines. GABA-producing neurons were targeted using *Gad1-GAL4* (*Gad1>*), which drives expression in neurons expressing the GABA-biosynthetic enzyme glutamate decarboxylase 1 (GAD 1). *Rdl-GAL4* (*Rdl>*) was used to drive expression in neurons expressing the ionotropic GABAA receptor, while *GABA-B-R2-GAL4* (*GABA-B-R2>*) was used to target neurons expressing the metabotropic GABAB receptor subtype 2. Furthermore, GABAergic innervation has been shown to regulate “clock” neurons expressing the Pigment Dispersing Factor (PDF) peptide, comprising a limited number of wake-promoting neurons, the small and large ventral lateral neurons (s-LNv and l-LNv, respectively) [40, 41]. To assess the function of *nowl* in these neurons, the *Pdf-GAL4* (*Pdf>*) driver was used to drive expression in all PDF-expressing neurons, while the *dimm-GAL4* (*dimm>*) driver allowed for knockdown in peptidergic neurosecretory cells including the l-LNvs, suggested to be the only sleep-regulatory neurons in this subset of cells [42]. Since global or neuronal loss of *nowl* leads to highly fragmented night-time sleep, we asked whether knockdown of *nowl* in any of these restricted neuronal subsets could elicit a similar effect on sleep maintenance. Interestingly, we found that knockdown of *nowl* using the *Rdl>* driver, thus specifically in GABAA-receptor*-*expressing neurons, caused a shortening of sleep bouts and an increase in their number during night, with an effect size similar to that seen with pan-neuronal knockdown (Fig. 7A-7B). *Rdl* encodes the *Drosophila* GABAA receptor, a ligand-gated chloride channel that mediates fast inhibitory effects of GABA, and it has been shown to affect several behaviors such as learning and sleep [39, 43]. Knockdown of *nowl* in other neuronal populations using the other drivers – targeting the GABA-producing cells, the GABAB-receptor-expressing neurons, and the PDF-expressing clock neurons – had no significant effect on night-time sleep-bout number or duration.

**Figure 7.**
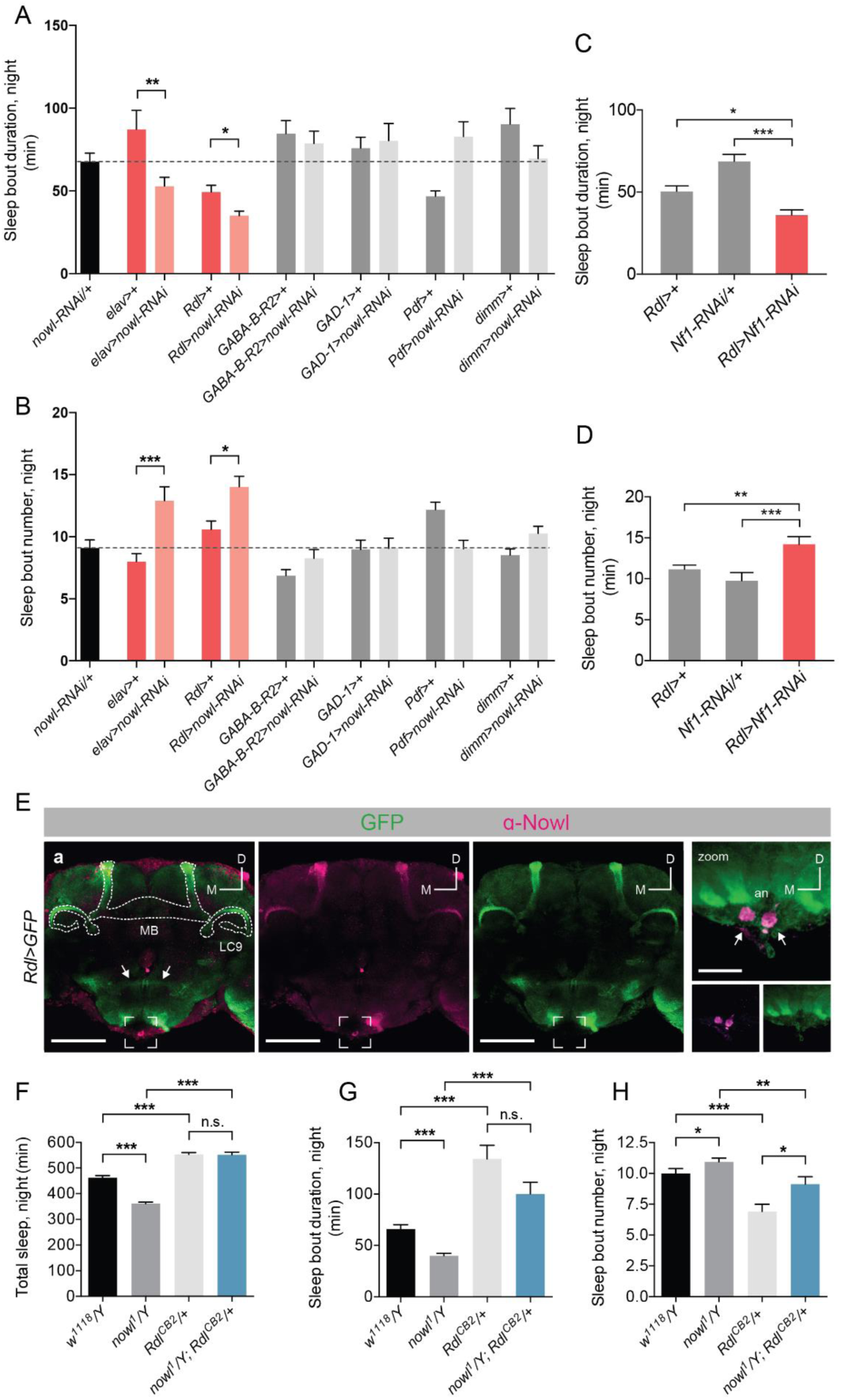
*nowl* is required in GABA-responsive neurons for sleep maintenance, and increased GABA signaling rescues the sleep phenotype of *nowl* mutants. (A-D) Quantification of average male night-time sleep-bout duration and length. (A) Sleep-bout number is increased, and (B) sleep-bout duration is reduced during the night when *nowl* is knocked down pan-neuronally (*elav>nowl-RNAi*) or specifically in GABAA-receptor *Rdl*-expressing neurons (*Rdl>nowl-RNAi*) compared to controls (*elav>*+ and *Rdl>+*), respectively. Knockdown animals were compared to the *GAL4* driver line crossed to *w*^*1118*^ (the genetic background for the RNAi), and *elav>nowl-RNAi* was used as a positive control. (C) Sleep-bout duration is decreased and (D) sleep-bout number increased in animals with knockdown of *Nf1* in *Rdl*-expressing neurons compared to controls (*Rdl>+* and *Nf1-RNAi/+*). (E) Anti-Nowl staining (purple) is prominent in *Rdl*-expressing neurons (*Rdl>GFP*, green) of the mushroom body (MB), among other sites. The neurons of the α′/β′ lobes of the MB (outlined; “MB”) and the neurons projecting to the LC9 optic glomerulus (outlined; “LC9”) express both *Rdl>GFP* and Nowl. Interneurons connecting the paired olfactory lobes (arrows) express *Rdl>GFP* but not Nowl, and a pair of neuronal clusters at the tip of the subesophageal ganglion (bracketed and zoomed in right panel) strongly express Nowl but not *Rdl>GFP*, indicating independent expression and staining. (F-H) Quantification of male night-time total sleep and average sleep-bout duration and number for *w*^*1118*^*/Y, nowl*^*1*^*/Y, Rdl*^*CB2*^*/+*, and *nowl*^*1*^*/Y; Rdl*^*CB2*^*/+*. (F) The reduced night-time sleep exhibited by *nowl*^*1*^ mutant flies is rescued by introducing one copy of the *Rdl*^*CB2*^ allele. (G, H) Sleep-bout duration during the night is significantly decreased in *nowl* mutants, and sleep-bout number is significantly increased. Introducing one copy of the *Rdl*^*CB2*^ allele partially rescues both defects. Graphs represent means with SEM (n=32-156). For normally distributed data, significance was determined using a one-way ANOVA with Dunnetts post-hoc testing. For non-normally distributed data, Kruskal-Wallis test with Dunns post-hoc testing was used (* p<0.05, ** p<0.01, *** p<0.001).

Since our data show that *nowl* interacts genetically with *Nf1* in the regulation of night-time sleep (Fig. 4 and 5), we asked whether *Nf1* also regulates sleep maintenance through specific effects in the GABAA-receptor Rdl-expressing neurons, like *nowl*. Consistent with the interaction of *nowl* and *Nf1* in sleep regulation, knockdown of *Nf1* in *Rdl*-expressing neurons produced a sleep fragmentation phenotype characterized by increased sleep bouts and decreased duration similar to loss of *nowl* in these GABA-responsive neurons (Fig. 7C-7D). Together these data suggest that *nowl* and *Nf1* are specifically required in GABA-responsive *Rdl*-expressing neurons to maintain sleep after sleep onset. The lack of knockdown effect in the PDF-expressing neurons further supports the independence of the circadian clock from *nowl* function.

To investigate the expression of Nowl in the brain, particularly in GABAA-receptor *Rdl-*expressing neurons, we generated an anti-Nowl antibody. Colocalization of mCD8::GFP expression in the *Rdl*-expressing neurons with staining against Nowl shows that the mushroom bodies (MBs) express both the GABAA-receptor *Rdl* and Nowl (Fig. 7E). Interestingly, downregulation of *Rdl* in the α′/β′ neurons of the MBs causes sleep loss similar to knockdown of *nowl* [44]. Together these data are consistent with a model in which Nowl promotes night-time sleep via modulation of GABA signaling that regulates the activity of the Rdl-expressing mushroom-body α′/β′ lobes.

### Increased inhibitory signaling through the GABAA receptor rescues *nowl* mutant sleep phenotype

Knockdown of *nowl* in GABAA-receptor *Rdl*-expressing neurons led to disruptions in sleep architecture, similar to the effects observed when decreasing inhibitory GABAergic signaling via Rdl. We therefore hypothesized that loss of *nowl* might affect sleep by reducing inhibitory GABAergic signaling effects in these neurons, thus disinhibiting them, which should promote wakefulness and reduce sleep [38, 39]. To address this possibility, we asked whether the *Rdl*^*CB2*^ mutation would rescue the *nowl* mutant sleep phenotype. *Rdl*^*CB2*^ mutant flies express a form of the GABAA receptor Rdl that is thought to exhibit reduced desensitization, leading to longer channel-opening duration, increased channel flux, and therefore increased neuronal inhibition downstream of GABA reception [37]. We rationalized that if loss of *nowl* increases neuronal excitability due to loss of GABAA-receptor signaling in *Rdl*-expressing neurons, the mutant Rdl^CB2^ receptor might compensate for this and thereby rescue the sleep disturbance observed in *nowl* loss-of-function flies. Compared to control flies, introduction of one copy of the *Rdl*^*CB2*^ allele significantly increased total sleep during the night, consistent with the expected effect of increased inhibitory GABA signaling. As we previously observed, *nowl* mutant flies displayed significantly decreased total sleep during the night. This effect was completely rescued in flies carrying both *nowl*^*1*^ and *Rdl*^*CB2*^ (Fig. 7F). The addition of *Rdl*^*CB2*^ caused a decreased number of longer night-time sleep bouts, eliminating the night-time sleep fragmentation induced by the loss of *nowl* function (Fig. 7G-7H). Taken together, these data suggest that increased inhibitory signaling through the GABAA-receptor Rdl largely rescues the sleep disturbances observed in *nowl* loss-of-function flies. We thus propose that *nowl* affects sleep by regulating the excitability of GABA-responsive *Rdl*-expressing neurons.

## Discussion

Schizophrenia is a highly heritable disorder with a prevalence of 1% in the population [45, 46]. It encompasses a wide spectrum of symptoms, of which a major one is sleep disturbance [9]. The human 22q11.2 chromosomal deletion spans more than 40 genes, and children carrying this deletion are 20 to 25 times more prone to developing schizophrenia during adolescence [47]. To identify which 22q11.2 genes might underlie the syndrome and development of schizophrenia in human carriers, we screened for conserved genes within this deletion that are required for sleep in *Drosophila* by RNAi-induced knockdown in the nervous system. Neuronal knockdown of several genes each caused abnormal sleep patterns and activity in flies. In females, silencing of several genes including *CDC45L* (human *CDC45*), *CG13192* (*GNB1L*), *CG13248* (*SLC7A4*), *Crk (CRK), Es2* (*DGCR14*), *Septin4* (*SEPT5*), and *UFD1-like* (*UFD1L*) resulted in short-sleep phenotypes. Some of these genes have not previously been linked to behavioral deficits in schizophrenia. However, we observed that neuronal knockdown of *CG13192* (human *GNB1L*) and *slgA* (*PRODH*) resulted in an increased number of sleep episodes that have a shorter length. PRODH catalyzes a step in the conversion of proline to glutamate, while *GNB1L* encodes a G-protein beta-subunit-like polypeptide [48, 49]; both genes have also been associated with schizophrenia and other mental disorders [48, 50-52]. In males, neuronal knockdown of several genes increased total sleep, while only *nowl* knockdown caused a decrease in total sleep. We show here that *nowl*, the conserved *Drosophila* homolog of the human *LZTR1* gene contained within the 22q11.2 deletion, is required for night-time sleep initiation and maintenance. Mutation or neuronal knockdown of *nowl* decreased total sleep amount and caused highly fragmented sleep, especially during night-time, which is a phenotype frequently seen in mental illness [26]. In addition, *nowl* mutants exhibit highly delayed sleep onset (*i.e.*, increased sleep-onset latency). Sleep-onset latency is a widely used measure in the study of human sleep disorders and psychiatric illnesses, including schizophrenia [26]. Accumulating evidence also indicates that getting enough sleep is important for the maintenance of energy balance and that reduced or poor sleep increases the risk of developing obesity [31]. Sleep is tightly connected to metabolic processes, and one proposed function of sleep is that it serves a key role in replenishing glycogen stores, which are depleted during wakefulness [32, 33]. Consistent with this notion, brain glycogen levels are highest during periods of sleep and decrease following rest deprivation in *Drosophila* [53]. Our findings of reduced glycogen levels in flies lacking *nowl* and *Nf1* function agrees with the view that sleep disruption is associated with depletion of glycogen stores, although further studies are needed to determine whether this effect is linked to alteration in sleep or more direct effects on metabolism. Furthermore, our results show increased adiposity associated with loss of *nowl* in flies, suggesting that *LZTR1* loss may contribute to the increased genetic susceptibility to develop obesity found in human 22q11.2 carriers [34]. Taken together the defects in sleep initiation and maintenance observed in *nowl* mutants suggest that *LZTR1* loss may contribute to the symptoms observed in the 22q11.2 DS and in schizophrenia, including the increased incidence of obesity.

Delayed sleep onset and the short-sleep phenotype could be caused either by alterations in sleep homeostasis or by disturbances in circadian rhythm. Our data indicate that animals lacking *nowl* function exhibit a normal circadian rhythm, which suggests that their altered sleep is not related to problems with circadian sleep regulation. Furthermore, the night-time short sleep of animals with loss of *nowl* function is not a consequence of hyperactivity, which demonstrates that the sleep fragmentation observed in these animals is a specific sleep-disturbance phenotype. Since *nowl* encodes a highly conserved protein (51% identity and 67% similarity to the human LZTR1 protein), Nowl may govern an evolutionarily conserved mechanism that regulates night-time sleep in animals.

Human LZTR1 has been shown to interact with Cul3 in the Cul3 ubiquitin-ligase complex [27]. Cul3 in *Drosophila* has been associated with sleep and interacts physically in this regulation with Insomniac, another BTB/POZ-domain protein (like Nowl) that functions as a substrate adaptor in the Cul3 complex [15, 30]. Although *Cul3* knockdown and *nowl* knockdown are phenotypically similar, *Cul3* knockdown induces broader phenotypes including disruption of the circadian clock [14]. Future studies should determine whether Nowl regulates sleep through a physical interaction with Cul3.

Recent studies have found that LZTR1 may function in the Cul3 ubiquitin ligase complex to ubiquitinate Ras [27, 28], which has previously been implicated in sleep regulation [54]. Nf1 is an important regulator of Ras signaling, and mutations in *Nf1* and *LZTR1* both give rise to types of neurofibromatosis [17, 55]. We show that *nowl* is required for proper negative regulation of Ras signaling, similar to Nf1, and that *nowl* and *Nf1* interact in the regulation of sleep. Heterozygotes for either *nowl* and *Nf1* mutations exhibit normal sleep patterns, while *nowl Nf1* trans-heterozygous animals display disrupted sleep architecture, including sleep fragmentation, indicating a genetic interaction. Furthermore, the reduction in total sleep and the fragmentation of night-time sleep caused by loss of *nowl* are partially rescued by *Nf1* overexpression. Thus, these two genes may function in the same pathway, or both of them may act on a common element involved in sleep maintenance. Whether *nowl* and *Nf1* regulate sleep through their effects on Ras activity will be an interesting question for future studies.

GABA neurotransmission is often altered in schizophrenic patients, and it is the main system that regulates night-time sleep [37, 41, 56, 57]. Since disruption of GABA signaling has also been shown to reduce night-time sleep in *Drosophila*, we asked whether *nowl* and *Nf1* might function in GABA-producing or -responsive cells. We found that knockdown of *nowl* in *Rdl*-expressing GABA-responsive neurons led to strongly increased sleep fragmentation, similar to the effect of pan-neuronal *nowl* knockdown. The wake-promoting l-LNv “clock” neurons express Rdl and play a major role in regulating sleep [39]. When these neurons are artificially hyper-excited by the expression of the sodium channel NaChBac, which allows membrane depolarization to occur more readily, flies showed decreased levels of sleep and increased levels of arousal, especially during night-time [42]. Knockdown of *Rdl* in these cells increases sleep latency [38], while reduced *GABAB-R-2* metabotropic receptor expression leads to reduced sleep during the late night [41]. Using RNAi-induced knockdown driven by the *Pdf>* and *dimm>* constructs, we found that *nowl* does not appear to function in these s-LNv clock neurons to regulate sleep, indicating that other *Rdl*-expressing neurons are involved in sleep-regulatory *nowl* activity. The mushroom bodies (MBs), which are required for olfactory associative learning and memory [58] and which also regulate sleep [59, 60], strongly express Rdl (Fig. 7E and [43]). A single pair of dorsal paired medial (DPM) neurons have been found to strongly promote sleep via GABAergic and serotonergic innervation of the mushroom bodies [44]. The MB α′/β′ neuronal population in particular are wake-promoting neurons that receive GABAergic input from the DPM neurons. Increased activation of the α′/β′ population decreases night-time sleep and increases sleep fragmentation [44]. Knockdown of *Rdl* in the α′/β′ neurons results in night-time sleep-loss phenotypes almost identical to those seen with knockdown of *nowl* pan-neuronally or specifically in the *Rdl*-expressing neurons, inducing increased sleep-bout numbers and reduced bout lengths, with a stronger effect during night-time [44]. Furthermore, we found strong enrichment of Nowl in *Rdl*-expressing MB neurons, consistent with Nowl’s functioning in the regulation of sleep via wake-promoting neurons in the mushroom body. Interestingly, *nowl* was also identified in an olfactory-learning screen, in which it was found to inhibit electric-shock-reinforced associative learning, consistent with a role for *nowl* in mushroom-body neuronal function [25]. Furthermore, we observed that *Nf1* knockdown in *Rdl*-expressing neurons caused sleep reduction and fragmentation of night-time sleep similar to *nowl* knockdown, further supporting an interaction between *Nf1* and *nowl* in the regulation of sleep. Investigating whether *nowl* and *Nf1* do indeed regulate sleep via effects on GABA signaling in the mushroom bodies will be an interesting subject for follow-up studies.

It has been suggested that mutations in ion channels can lead to disrupted sleep by altering overall neuronal excitability [61]. One example of this is the gene *Shaker*, which encodes an α-subunit of a potassium channel that functions to regulate membrane repolarization after neuronal depolarization [62]. Loss-of-function mutations in this gene, and in its mouse orthologs, cause reduced sleep and shorter sleep episodes, without affecting circadian or homeostatic sleep drive [63]. Since sleep loss and fragmentation due to loss of *nowl* were partially rescued by increased GABAergic inhibition of *Rdl*-expressing neurons, we suggest that loss of *nowl* may alter neuronal excitability in wake-promoting *Rdl*-expressing neurons. It will be interesting to determine whether *nowl* directly affects the Rdl receptor or other substrates via Cul3 to regulate postsynaptic GABA signaling. Cul3 and its other interaction partner Insomniac are recruited to the postsynaptic compartment within minutes of acute glutamate-receptor inhibition [64]. These proteins mediate local mono-ubiquitination, which is important for homeostatic signaling in the postsynaptic compartment. The number of Rdl receptors expressed in a neuron, beyond the mere presence or absence of these receptors, has been suggested to be important in sleep regulation [39]. Furthermore, in humans, agonists of GABAA-type receptors are common treatments for insomnia [65], and a loss-of-function mutation affecting a GABAA receptor subunit causes a heritable type of insomnia [66]. Altered GABAergic signaling is believed to play a role in many neurodevelopmental disorders, and a mechanism that involves GABAA-receptor signaling may be a key factor underlying the pathophysiology of the 22q11.2 DS. Our findings suggest a contribution of *nowl/LZTR1* – a 22q11.2 gene – in altered GABAergic neurotransmission and thus may provide a new direction for understanding the mechanisms underlying the observed neurodevelopmental phenotypes in the 22q11.2 DS and for developing therapeutic interventions against them.

Essentially all psychiatric disorders are associated with sleep disturbances. We have identified *nowl* as regulator of sleep in *Drosophila*. Mutations in its human ortholog *LZTR1* have been implicated in several human diseases, such as schwannomatosis, glioblastoma, and the psychiatric disorders linked to the 22q11.2 DS. We suggest that the *nowl/LZTR1* gene encodes a conserved regulator of sleep that interacts with Nf1 and that may contribute to the overall pathophysiology of the 22q11.2 DS. Further studies of Nowl may help understand the molecular mechanisms underlying both mental disorders and Schwann-cell tumors.

## Materials and methods

### *Drosophila* lines and maintenance

*Drosophila* larvae and adults of mixed sexes were raised on standard cornmeal medium (Nutri-Fly “Bloomington” formulation) at 25 °C under a 12:12 light/dark cycle with 60% humidity. The following fly lines were obtained from Bloomington *Drosophila* Stock Center (BDSC; Bloomington, IL): *elav-GAL4; UAS-Dicer-2* (#25750), *nSyb-GAL4* (#51635), *Gad1-GAL4/CyO* (#51630), *Rdl-GAL4/CyO* (#66509), *Pdf-GAL4* (#6899), *dimm-GAL4* (#25373), *UAS-Dicer-2* (#24651), *Mi{ET1}CG3711*^*MB12128*^ (#29940), *Rdl*^*CB2*^*/TM6B, Tb*^*1*^ (#35493), and *UAS-mCD8::GFP* (#4776). *UAS-RNAi* lines against orthologs of human 22q11.2 CNV-linked genes were procured from Vienna *Drosophila* Resource Center (VDRC; Vienna, Austria) and are listed in Supplementary Table 1. *UAS-Cul3-RNAi* (#109415), *UAS-Nf1-RNAi* (#109637), and the *w*^*1118*^ (#60000) genetic background line were also obtained from VDRC.

The *GABA-B-R2*-*GAL4*::*p65* line carries a bacterial artificial chromosome (BAC) containing ∼80 kb of genomic sequence surrounding the *GABA-B-R2* gene, in which the first coding exon of this gene was replaced with *GAL4::p65* and associated terminator sequences; the remaining exons and introns are still present, with the assumption that they contain regulatory sequences, but they are no longer transcribed. This construct was built using recombineering techniques [67] in P[acman] BAC clone CH321-95N09 (Children’s Hospital Oakland Research Institute, Oakland, CA). Selectable markers (conferring kanamycin resistance and streptomycin sensitivity) were cloned from *pSK+KanaRpsL* [68] (AddGene plasmid #20871) and flanked with upstream *GAL4* and downstream *HSP70-UTR* arms cloned from *pBPGUw* [69], a gift of G. Rubin. *GABA-B-R2*-specific homology arms were added by PCRing this landing-site cassette with the following primers (genomic sequence is in lower case): GABA-B-R2-GAL4-F: cgatatgcgc tattcacatt tagaatcgtt ttacagccca cgcggtcaac ATGAAGCTAC TGTCTTCTAT CGAACAAGC; GABA-B-R2-UTR-R: catcatcaga gattcactta atgaaatctt caagctaaac cctaactcac GATCTAAACG AGTTTTTAAG CAAACTCACT CCC. The homology-flanked landing-site cassette was recombined into the BAC (with kanamycin selection), and then full-length *GAL4*::*p65-HSP70* amplified from *pBPGAL4.2::p65Uw* [70] (a gift of G. Rubin) was recombined into this site in a second recombination (with streptomycin selection). The recombined regions of the BAC were sequenced, and the sequence-verified BAC was integrated into the *Drosophila* genome at *attP40* [71] by Genetic Services, Inc. (Cambridge, MA).

### Identification of *Drosophila* orthologs

*Drosophila* orthologues of human 22q11.2 CNV-linked genes were identified using the “*Drosophila* RNAi Screening Center Integrative Ortholog Prediction Tool” (DIOPT), available at the website of the *Drosophila* RNAi Screening Center (DRSC), Harvard Medical School [21].

### Sleep assays and analysis

The *Drosophila* Activity Monitor (DAM) system (TriKinetics, Inc., Waltham, MA) was used to measure locomotion over a 24-hour period. Adult flies were sorted by sex and collected in groups of 30 soon after eclosion. Collected flies were housed under standard conditions until the start of the experiment. Three-to-seven-day-old flies (either male only or female only, as indicated) were used for experiments, in which they were housed in 65-mm-long glass tubes containing a plug of food medium (5% sucrose and 2% agar in water) at one end and a cotton stopper at the other. Experiments were run in a behavioral incubator under a 12-hour light/12-hour dark cycle, and flies were allowed 12-24 hours to acclimatize prior to experimental start. Activity, determined as the number of beam crosses per fly per minute, was measured in one-minute periods, and episodes of sleep were defined as at periods of least 5 minutes of uninterrupted quiescence. Flies exhibiting less than 10 minutes of activity during either the entire light or dark phase were flagged as dead. The pySolo software package [72] and a custom MATLAB script (MATLAB R2016b, The MathWorks Inc, Natick, Massachusetts) written by Stanislav Nagy were used to analyze sleep dynamics. Sleep latency was calculated as the period from lights-off to the first sleep episode. Locomotor activity of male flies kept on a normal LD cycle for several days was monitored in constant darkness (DD) for 8 days to determine the length of the free-running circadian period. Data were aggregated into 30-minute bins and analyzed using the FaasX software [73], which uses Chi-Square test to calculate the period length.

### Western blotting

For Western-blot analysis, ten male flies per genotype were frozen at -80 °C, vortexed, and sieved to obtain samples of heads. Heads were homogenized in 50 μl Laemmli Sample Buffer (Bio-Rad) containing 2-mercaptoethanol. Samples were denatured for 5 minutes at 95 °C and then centrifuged at maximum speed (17,000 g) for 5 minutes. Samples (20 μl) were loaded on 4-20% gradient polyacrylamide gel (Bio-Rad), and proteins were separated at 150 V for 30 minutes. Proteins were transferred onto PVDF membrane (Millipore), and the membrane was blocked for one hour in Odyssey Blocking Buffer (LI-COR). Primary antibodies (rabbit anti-phospho-ERK, Cell Signaling Technology #9101, 1:1000; mouse anti-α-tubulin, Sigma-Aldrich #T9026, 1:5000) were diluted in Odyssey Blocking Buffer containing 0.2% Tween 20, and the membrane was incubated overnight in this solution. Membranes were then rinsed and incubated for 40 minutes with secondary antibodies conjugated to infrared fluorophores (anti-mouse IRDye 680RD and anti-rabbit 800CW, LI-COR, 1:10,000), and staining was imaged using an Odyssey Fc scanner (LI-COR).

### Anti-Nowl antiserum development and immunocytochemistry

To generate a Nowl antibody, a peptide corresponding to residues 123–141 of the parent protein (LRSSFKSSKRNKARKSAST) was used in a custom immunization protocol carried out by Genosphere Biotechnologies (Paris, France). Epitope specificity of the antiserum was confirmed by comparing wild-type (*Rdl>GFP*) and *nowl-RNAi* animals by immunostaining as well as by co-application of the pre-immune serum (S6C Fig.). For immunocytochemistry, primary antibodies used were polyclonal rabbit α-Nowl (1:200) and FITC-conjugated polyclonal goat anti-GFP (1:200; Abcam, #ab6662). Biotinylated anti-rabbit was applied (1:200; Vector Laboratories, #BA-1100), followed by Cy3-streptavidin (1:200; Jackson ImmunoResearch Laboratories, #016-160-084). Tissues were mounted on poly-L-lysine-coated 35-mm glass-bottomed dishes (MatTek Corporation, MA, USA) in Vectashield (Vector Laboratories Inc., CA, USA), and image acquisition was performed on an inverted Zeiss LSM800 confocal microscope with AiryScan (Zeiss, Oberkochen, Germany). Images were processed in CorelDraw X8 (Corel, Ottawa, Canada).

### Transcript analysis by quantitative PCR

The efficiency of *nowl* and *Nf1* RNAi-mediated gene knockdown was analyzed using quantitative RT-PCR (qPCR). Heads from adult male flies expressing neuronal RNAi against *nowl* or *Nf1*, along with driver- and RNAi-alone controls, were collected, and total RNA was extracted from 10 heads per sample using the RNeasy Mini Kit (Qiagen) according to the manufacturer’s instructions. cDNA was reverse transcribed from total RNA using the High-Capacity cDNA Reverse Transcription Kit (ThermoFisher #4368814), and used a template for qPCR with the QuantiTect SYBR Green PCR Kit (Fisher Scientific #204145) and an Mx3005P qPCR System (Agilent Technologies). Levels of target gene expression was normalized against *RpL32*. Precomputed oligo pairs from the FlyPrimerBank was used for target gene primers. Primers were: GGAGGATCGGGATGTAGTATGG and CGTGTAGGAGGTGAAGTCCAC for *nowl*; AGTATCTGATGCCCAACATCG and CAATCTCCTTGCGCTTCTTG for *Nf1*; and CTTTTGGCACGTTTCGAGGAT and GGTAGCGCGATATGTGGATCAG for *RpL32*.

### Measurements of glycogen and triglycerides

For triglyceride and glycogen assays, one-week old adult males flies were collected and frozen at -80 °C in groups of three animals until assayed. Animals were homogenized in 80 µl PBS + 0.5% Tween using a Tissue Lyser LT (Qiagen) with small steel beads. For triglyceride measurements, 8 µl of the homogenate was added to 80 µl Free Glycerol Reagent (Sigma, F6428) and 20 µl Triglyceride Reagent (Sigma, T2449), and reactions were incubated at 37 ^°^C for 10 minutes with agitation, before being read at 540 nm absorbance. For glycogen measurements, 6 µl homogenate was mixed with 94 µl Glucose Oxidase (GO) reagent from the Glucose (GO) Assay Kit (Sigma, GAGO20) containing 1 µl (0.3 U) of amyloglucosidase (Sigma, A7420-5MG) to determine total glucose + glycogen. Another 6 µl homogenate was mixed with 94 µl GO reagent alone (without the amyloglucosidase) to determine free glucose. Reactions were incubated at 37 ^°^C for 20 minutes, after which 66 µl of 12-N sulfuric acid was added to stop and complete the reaction before absorbance was measured at 540 nm. Absorbance was measured using an EnSight multimode plate reader (PerkinElmer), and values were standardized against readings for a range of known concentrations.

### Statistics

GraphPad Prism software was used for statistical analysis. Data was tested for normality using D’Agostino-Pearson normality test. One-way ANOVA with Dunnett’s multiple comparisons was performed to establish differences between control and several genotypes for normally distributed data, while Kruskal-Wallis test with Dunn’s post-hoc testing was used for non-normally distributed data. Statistical significance was set at p≤0.05.

## Supplementary materials

**S1 Fig.**
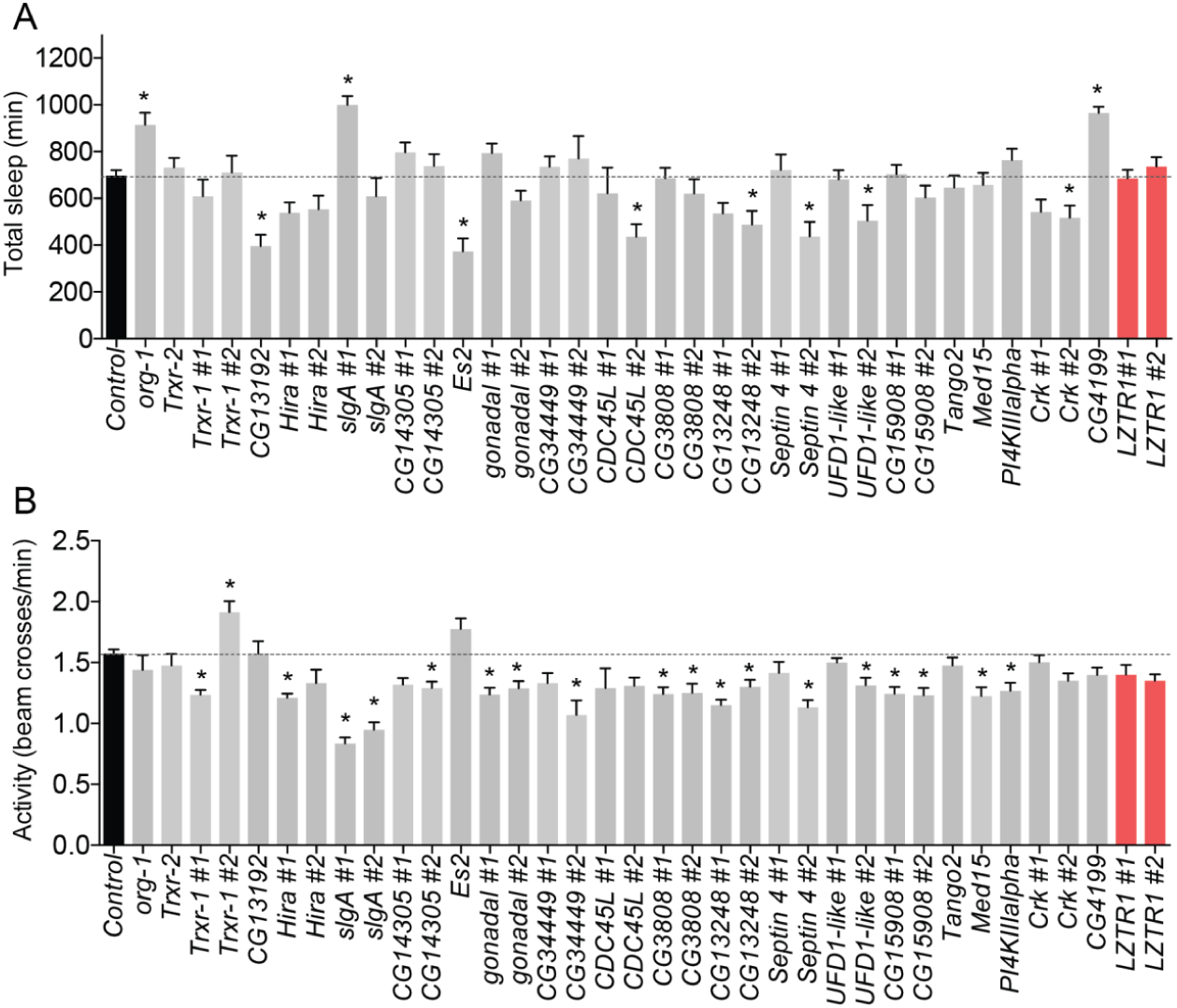
Screening individual *Drosophila* orthologs of genes spanned by the human 22q11.2 deletion for effects on sleep and activity in adult females. (A, B) Quantification of total sleep (A) and average activity measured as the number of beam crosses per minute (B) in 3-to-7-day-old females, measured over a 24-hour period. *UAS-RNAi* constructs were expressed under the control of the pan-neuronal *elav-GAL4* driver. n=140 flies for controls (*elav>* crossed to *w*^*1118*^, the genetic background for the RNAi lines); n=16 flies for each RNAi genotype. When possible, two distinct *UAS-RNAi* constructs were used against each gene. RNAi efficiency was enhanced by co-expressing the processing enzyme Dicer-2 (*UAS-Dcr-2*). Graphs represent means with SEM. Statistical significance was determined using a one-way ANOVA with Dunnetts post-hoc testing (* p<0.05).

**S2 Fig.**
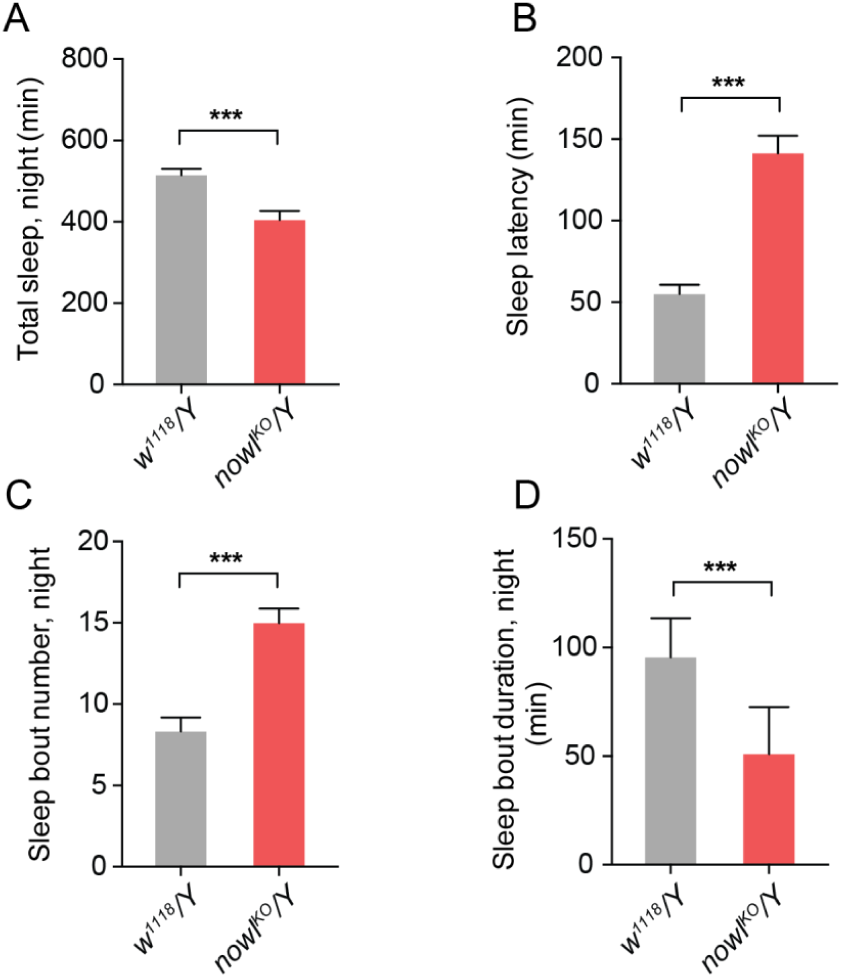
The CRISPR-induced *nowl*^*KO*^ mutation causes a sleep phenotype similar to that seen with the *nowl*^*1*^ insertion allele. (A, B) Total sleep per night (A) is decreased and sleep latency (B) is increased in *nowl*^*KO*^ mutants (*nowl*^*KO*^/Y) compared to the control (*w*^*1118*^/Y). (C, D) Night-time sleep is fragmented in *nowl*^*KO*^ mutants compared to the control, as indicated by increased sleep-bout number (C) and reduced sleep-bout duration (D). Graphs represent mean and SEM (n=32). Significance was tested by Mann Whitney test (*** p < 0.001).

**S3 Fig.**
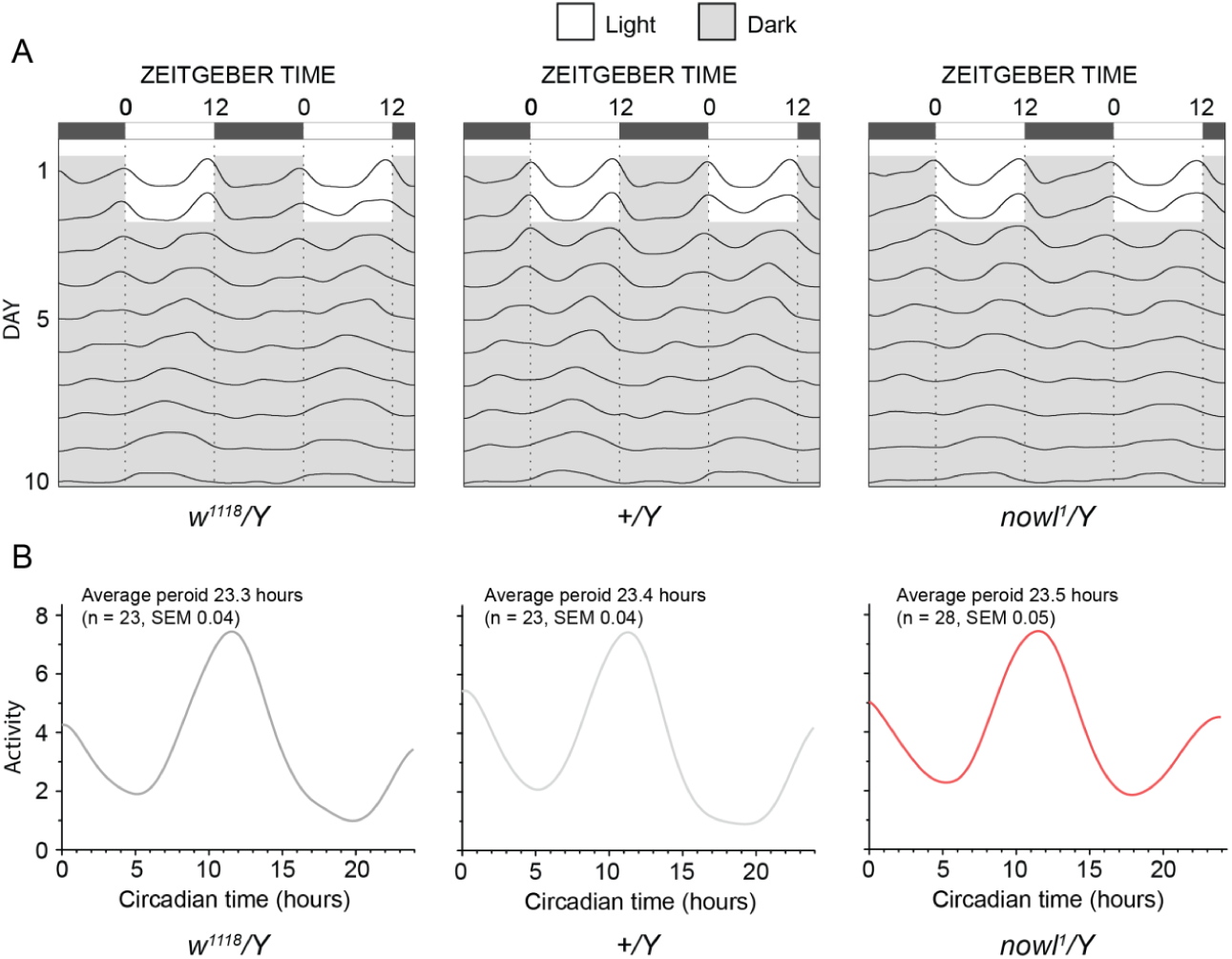
*nowl* mutants exhibit normal circadian rhythm. (A, B) Circadian rhythmicity of 3-to-8**-**day-old male flies kept in the dark for 8 days. During entrainment, lights-on occurred at Zeitgeber time (ZT) 0 and lights-off occurred at ZT12. (A) Actograms showing average activity during the light/dark entrainment period and during constant-darkness free-running cycles (ZT in hours). (B) Quantification of the free-running circadian period of *nowl*^*1*^ mutants indicates that it cycles similarly to controls (*w*^*1118*^/Y and +/Y) with a nearly 24-hour period, indicating that *nowl* is not necessary for entrainment or free running of the circadian clock. Data represents means (n=23-28). No significant differences by chi-square test in the FaasX software package [73].

**S4 Fig.**
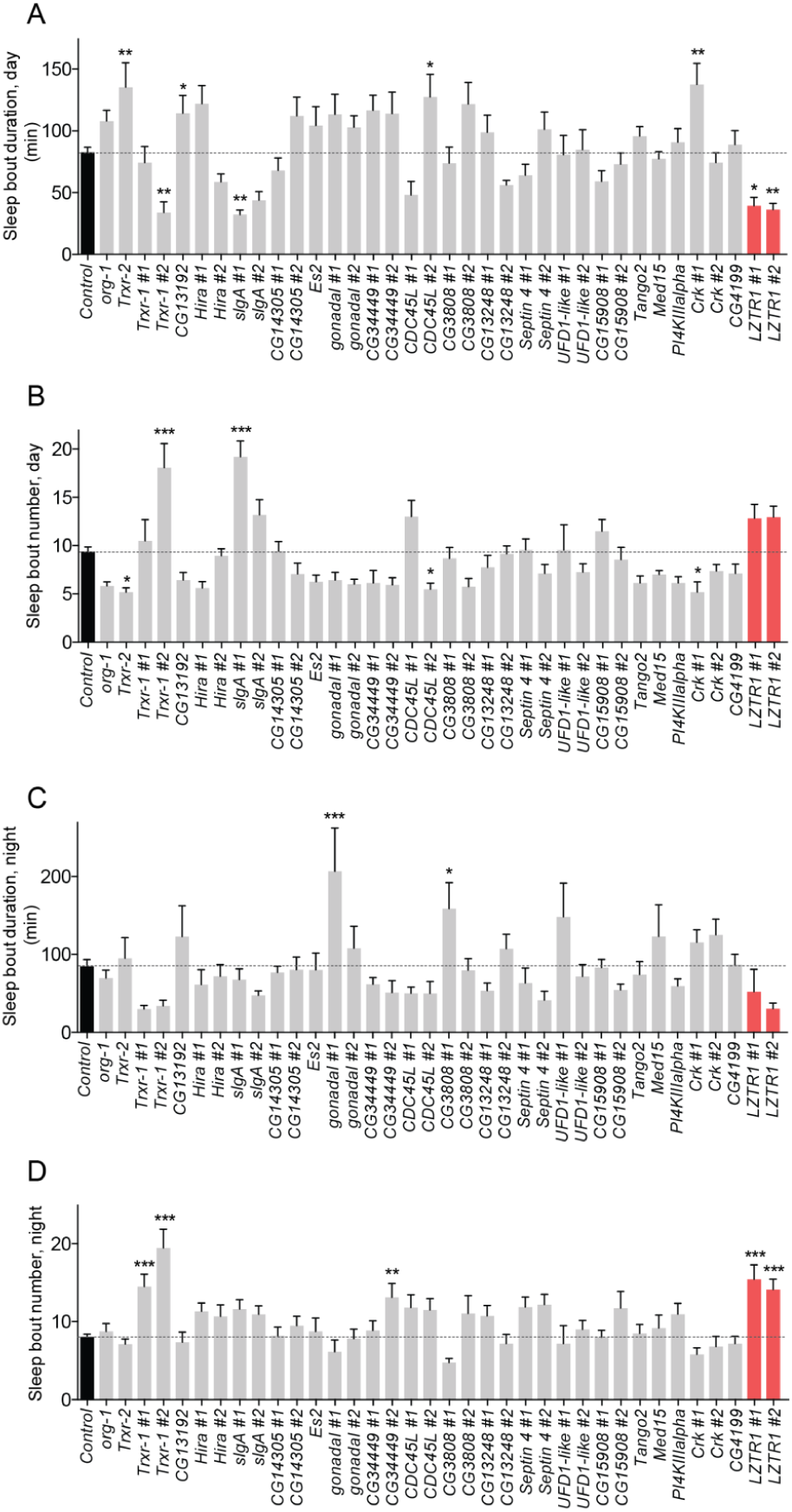
Screening individual *Drosophila* orthologs of human 22q11.2 deletion genes for effects on sleep maintenance. Neuronal gene knockdown was induced by crossing *elav>* to each gene-specific *UAS-RNAi* line. As a control, the *elav>* line was crossed to *w*^*1118*^. (A-D) Sleep-episode duration and number during the day (A, B) and night (C, D) of 3-to-8-day-old male flies (n=140 flies for controls; n=16 flies for each RNAi genotype). Graphs represent mean and SEM. Significance was tested by one-way ANOVA with Dunnett’s multiple-comparisons test (* p<0.05).

**S5 Fig.**
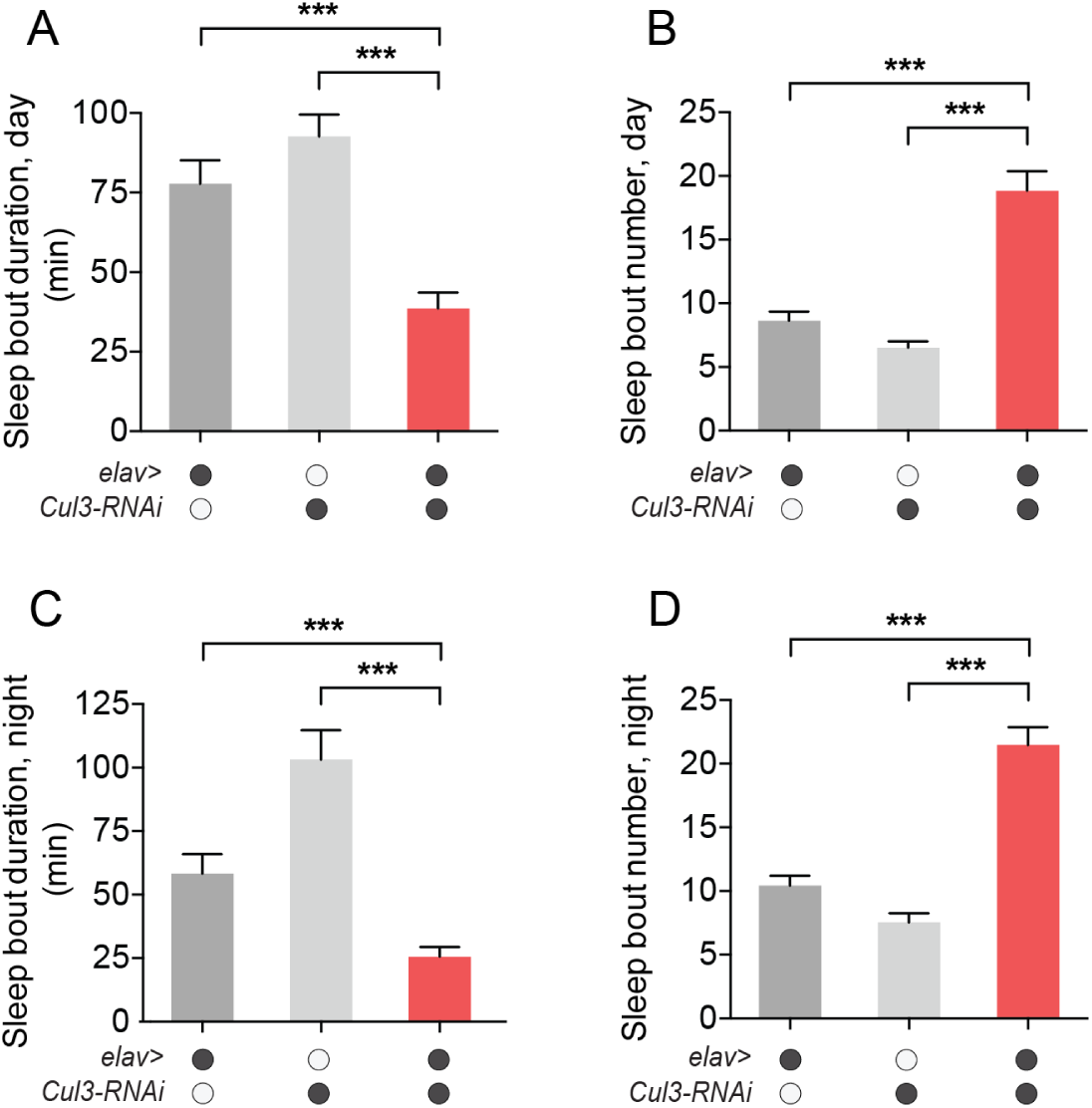
Pan-neuronal knockdown of *Cul3* causes sleep fragmentation. (A, B) Quantification of average male daytime sleep-bout duration (A) and number (B) for *elav>Cul3-RNAi* flies compared to controls (*elav>+* and *UAS-Cul3-RNAi/+*). Sleep-bout length was significantly decreased, and sleep-bout number was significantly increased during the day when *Cul3* was knocked down in the nervous system. (C, D) Quantification of average night-time sleep-bout duration (C) and number (D). Bout length was significantly decreased, and average bout numbers were significantly increased during the night upon pan-neuronal knockdown of *Cul3*. Graphs represent mean and SEM (n=32). Statistical significance was determined using Kruskal-Wallis test with Dunns post-hoc testing (*** p<0.001).

**S6 Fig.**
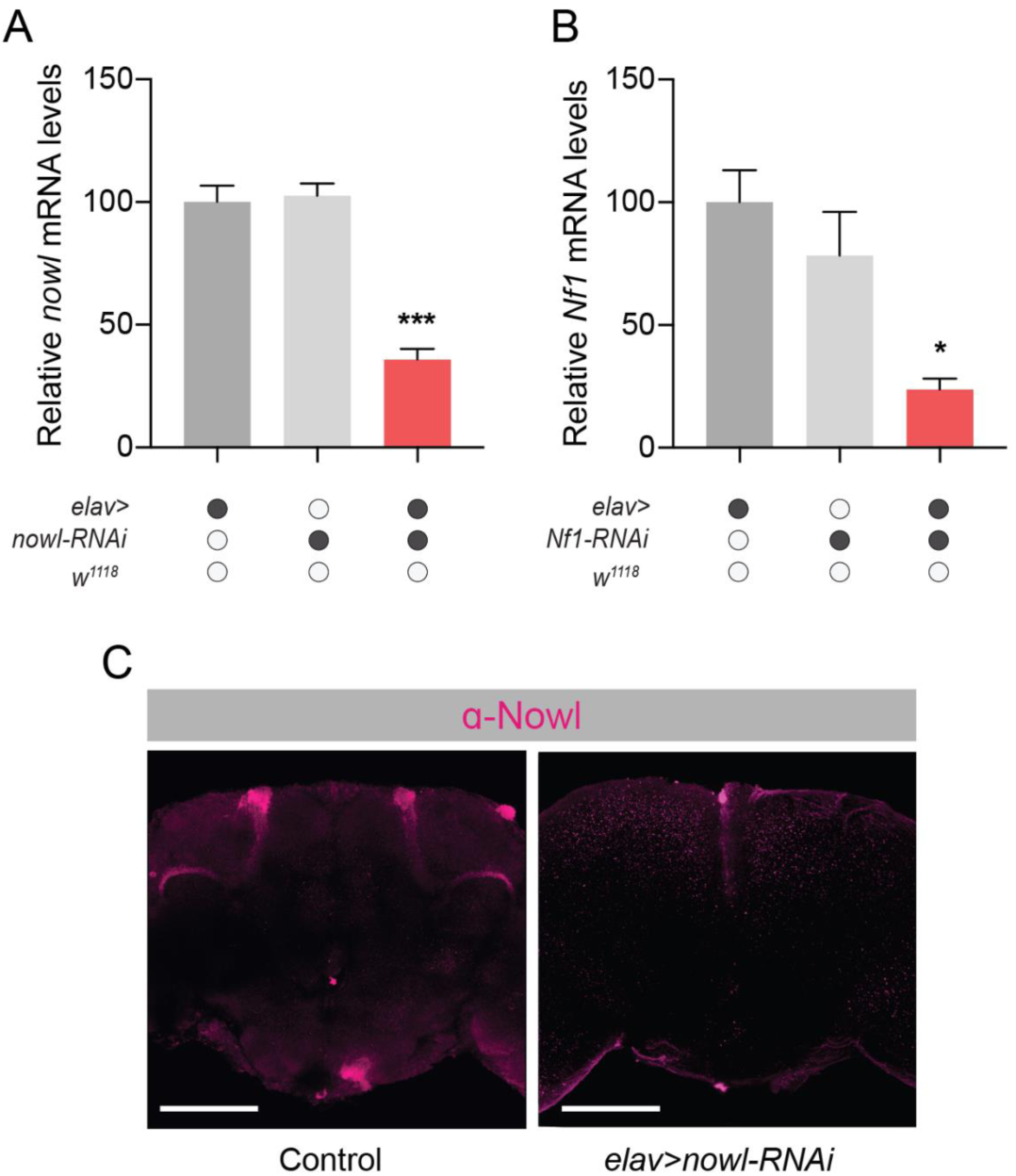
RNAi against *nowl* and *Nf1* is effective, and Nowl antibody staining is specific. (A, B) Efficiency of *nowl* (A) and *Nf1* (B) gene knockdown. Gene expression was determined in adult heads using qPCR. (C) Anti-Nowl staining features observed in *Rdl>GFP* controls are not present in *elav>nowl-RNAi* brains with reduced expression of *nowl* in the nervous system, indicating that the anti-Nowl antibody specifically recognizes the Nowl protein.

**Table S1.** Transgenic UAS-RNAi fly lines targeting orthologs of 22q11.2-linked genes.

| Human gene | *Drosophila* ortholog | Fly line (*UAS-RNAi*) from VDRC |
| --- | --- | --- |
| *PRODH* | *sluggish A (slgA)* | #101449<br>#18561 |
| *TSSK2* | *CG14305* | #17477<br>#17478 |
| *DGCR14* | *Es2* | #106243<br>#42996 |
| *SLC25A1* | *scheggia (sea)* | #109169<br>#50713 |
| *CLTCL1* | *Clathrin heavy chain (Chc)* | #103383<br>#23666 |
| *HIRA* | *Hira* | #106989<br>#13690 |
| *MRPL40* | *mRpL40* | #101442<br>#48166 |
| *C22orf39* | *CG15908* | #106340<br>#19621 |
| *UFD1L* | *Ubiquitin fusion-degradation 1-like (Ufd1-like)* | #104713<br>#24700 |
| *CDC45* | *CDC45L* | #41084<br>#20705 |
| *SEPT5* | *Septin 4 (Sep4)* | #109398<br>#7742 |
| *TBX1* | *Optomotor-blind-related-gene-1 (org-1)* | #104393<br>#37656 |
| *GNB1L* | *CG13192* | #110748<br>#32157 |
| *TXNRD2* | *Thioredoxin reductase-1, -2 (Trxr-1, -2)* | #16768<br>#50336 |
| *DGCR8* | *partner of drosha (pasha)* | #107445<br>#40118 |
| *TRMT2A* | *CG3808* | #108653<br>#34713 |
| *C22orf25* | *Transport and Golgi organization 2 (Tango2)* | #18175 |
| *ZDHHC8* | *Zinc finger DHHC-type containing 8 (Zdhhc8)* | #5520<br>#5521 |
| *DGCR6L* | *gonadal (gdl)* | #24453<br>#23456 |
| *MED15* | *Mediator complex subunit 15 (MED15)* | #21809 |
| *PI4KA* | *Phosphatidylinositol 4-kinase III $\alpha$ (PI4KIIIα)* | #105614<br>#15993 |
| *SNAP29* | *Synaptosomal-associated protein 29 kDa (Snap29)* | #107947<br>#18173 |
| *CRK* | *Crk oncogene (Crk)* | #106598<br>#19061 |
| *AIFM3* | *CG4199* | #106170<br>#26424 |
| *LZTR1* | *CG3711/Leucine zipper like transcription regulator 1 (Lztr1)* | #11164<br>#13008 |
| *SLC7A4* | *CG13248* | #102635 |

## References

1. Sebat J, Lakshmi B, Malhotra D, Troge J, Lese-Martin C, Walsh T, et al. Strong association of de novo copy number mutations with autism. Science (New York, NY). 2007;316(5823):445–9. Epub 2007/03/17. doi: 10.1126/science.1138659. PubMed PMID: 17363630; PubMed Central PMCID: PMCPMC2993504.

2. Stefansson H, Rujescu D, Cichon S, Pietilainen OP, Ingason A, Steinberg S, et al. Large recurrent microdeletions associated with schizophrenia. Nature. 2008;455(7210):232–6. Epub 2008/08/01. doi: 10.1038/nature07229. PubMed PMID: 18668039; PubMed Central PMCID: PMCPMC2687075.

3. Vo OK, McNeill A, Vogt KS. The psychosocial impact of 22q11 deletion syndrome on patients and families: A systematic review. American journal of medical genetics Part A. 2018;176(10):2215–25. Epub 2018/03/27. doi: 10.1002/ajmg.a.38673. PubMed PMID: 29575505; PubMed Central PMCID: PMCPMC6221171.

4. Malhotra D, Sebat J. CNVs: harbingers of a rare variant revolution in psychiatric genetics. Cell. 2012;148(6):1223–41. doi: 10.1016/j.cell.2012.02.039. PubMed PMID: 22424231; PubMed Central PMCID: PMC3351385.

5. Didriksen M, Fejgin K, Nilsson SR, Birknow MR, Grayton HM, Larsen PH, et al. Persistent gating deficit and increased sensitivity to NMDA receptor antagonism after puberty in a new mouse model of the human 22q11.2 microdeletion syndrome: a study in male mice. J Psychiatry Neurosci. 2017;42(1):48–58. PubMed PMID: 27391101; PubMed Central PMCID: PMCPMC5373712.

6. Shprintzen RJ. Velo-cardio-facial syndrome: 30 Years of study. Developmental disabilities research reviews. 2008;14(1):3–10. Epub 2008/07/19. doi: 10.1002/ddrr.2. PubMed PMID: 18636631; PubMed Central PMCID: PMCPMC2805186.

7. Szelenberger W, Soldatos C. Sleep disorders in psychiatric practice. World psychiatry: official journal of the World Psychiatric Association (WPA). 2005;4(3):186–90. PubMed PMID: 16633547.

8. Glickman G. Circadian rhythms and sleep in children with autism. Neurosci Biobehav Rev. 2010;34(5):755–68. doi: 10.1016/j.neubiorev.2009.11.017. PubMed PMID: 19963005.

9. Cohrs S. Sleep disturbances in patients with schizophrenia: impact and effect of antipsychotics. CNS drugs. 2008;22(11):939-62. PubMed PMID: 18840034.

10. Bassett AS, Chow EW, AbdelMalik P, Gheorghiu M, Husted J, Weksberg R. The schizophrenia phenotype in 22q11 deletion syndrome. The American journal of psychiatry. 2003;160(9):1580–6. Epub 2003/08/29. doi: 10.1176/appi.ajp.160.9.1580. PubMed PMID: 12944331; PubMed Central PMCID: PMCPMC3276594.

11. Yagi H, Furutani Y, Hamada H, Sasaki T, Asakawa S, Minoshima S, et al. Role of TBX1 in human del22q11.2 syndrome. Lancet (London, England). 2003;362(9393):1366–73. Epub 2003/10/31. PubMed PMID: 14585638.

12. Hendricks JC, Finn SM, Panckeri KA, Chavkin J, Williams JA, Sehgal A, et al. Rest in Drosophila is a sleep-like state. Neuron. 2000;25(1):129-38. PubMed PMID: 10707978.

13. Shaw PJ, Cirelli C, Greenspan RJ, Tononi G. Correlates of sleep and waking in Drosophila melanogaster. Science (New York, NY). 2000;287(5459):1834–7. Epub 2000/03/10. PubMed PMID: 10710313.

14. Grima B, Dognon A, Lamouroux A, Chelot E, Rouyer F. CULLIN-3 controls TIMELESS oscillations in the Drosophila circadian clock. PLoS biology. 2012;10(8):e1001367. Epub 2012/08/11. doi: 10.1371/journal.pbio.1001367. PubMed PMID: 22879814; PubMed Central PMCID: PMCPMC3413713.

15. Stavropoulos N, Young MW. insomniac and Cullin-3 regulate sleep and wakefulness in Drosophila. Neuron. 2011;72(6):964–76. Epub 2011/12/27. doi: 10.1016/j.neuron.2011.12.003. PubMed PMID: 22196332; PubMed Central PMCID: PMCPMC3244879.

16. Nagy S, Maurer GW, Hentze JL, Rose M, Werge TM, Rewitz K. AMPK signaling linked to the schizophrenia-associated 1q21.1 deletion is required for neuronal and sleep maintenance. PLoS Genet. 2018;14(12):e1007623. doi: 10.1371/journal.pgen.1007623. PubMed PMID: 30566533.

17. Williams VC, Lucas J, Babcock MA, Gutmann DH, Korf B, Maria BL. Neurofibromatosis type 1 revisited. Pediatrics. 2009;123(1):124–33. Epub 2009/01/02. doi: 10.1542/peds.2007-3204. PubMed PMID: 19117870.

18. Smith MJ, Isidor B, Beetz C, Williams SG, Bhaskar SS, Richer W, et al. Mutations in LZTR1 add to the complex heterogeneity of schwannomatosis. Neurology. 2015;84(2):141–7. doi: 10.1212/WNL.0000000000001129. PubMed PMID: 25480913.

19. Ballester R, Marchuk D, Boguski M, Saulino A, Letcher R, Wigler M, et al. The NF1 locus encodes a protein functionally related to mammalian GAP and yeast IRA proteins. Cell. 1990;63(4):851–9. Epub 1990/11/16. doi: 10.1016/0092-8674(90)90151-4. PubMed PMID: 2121371.

20. Hannan F, Ho I, Tong JJ, Zhu Y, Nurnberg P, Zhong Y. Effect of neurofibromatosis type I mutations on a novel pathway for adenylyl cyclase activation requiring neurofibromin and Ras. Hum Mol Genet. 2006;15(7):1087–98. Epub 2006/03/04. doi: 10.1093/hmg/ddl023. PubMed PMID: 16513807; PubMed Central PMCID: PMCPMC1866217.

21. Hu Y, Flockhart I, Vinayagam A, Bergwitz C, Berger B, Perrimon N, et al. An integrative approach to ortholog prediction for disease-focused and other functional studies. BMC bioinformatics. 2011;12:357. Epub 2011/09/02. doi: 10.1186/1471-2105-12-357. PubMed PMID: 21880147; PubMed Central PMCID: PMCPMC3179972.

22. Dietzl G, Chen D, Schnorrer F, Su KC, Barinova Y, Fellner M, et al. A genome-wide transgenic RNAi library for conditional gene inactivation in Drosophila. Nature. 2007;448(7150):151–6. Epub 2007/07/13. doi: nature05954 [pii] 10.1038/nature05954. PubMed PMID: 17625558.

23. Vissers JH, Manning SA, Kulkarni A, Harvey KF. A Drosophila RNAi library modulates Hippo pathway-dependent tissue growth. Nat Commun. 2016;7:10368. doi: 10.1038/ncomms10368. PubMed PMID: 26758424; PubMed Central PMCID: PMCPMC4735554.

24. Bellen HJ, Levis RW, Liao G, He Y, Carlson JW, Tsang G, et al. The BDGP gene disruption project: single transposon insertions associated with 40% of Drosophila genes. Genetics. 2004;167(2):761–81. Epub 2004/07/09. doi: 10.1534/genetics.104.026427. PubMed PMID: 15238527; PubMed Central PMCID: PMCPMC1470905.

25. Appel M, Scholz CJ, Muller T, Dittrich M, Konig C, Bockstaller M, et al. Genome-Wide Association Analyses Point to Candidate Genes for Electric Shock Avoidance in Drosophila melanogaster. PLoS One. 2015;10(5):e0126986. Epub 2015/05/21. doi: 10.1371/journal.pone.0126986. PubMed PMID: 25992709; PubMed Central PMCID: PMCPMC4436303.

26. Anderson KN, Bradley AJ. Sleep disturbance in mental health problems and neurodegenerative disease. Nat Sci Sleep. 2013;5:61–75. Epub 2013/06/14. doi: 10.2147/NSS.S34842. PubMed PMID: 23761983; PubMed Central PMCID: PMCPMC3674021.

27. Steklov M, Pandolfi S, Baietti MF, Batiuk A, Carai P, Najm P, et al. Mutations in LZTR1 drive human disease by dysregulating RAS ubiquitination. Science (New York, NY). 2018;362(6419):1177–82. Epub 2018/11/18. doi: 10.1126/science.aap7607. PubMed PMID: 30442762.

28. Bigenzahn JW, Collu GM, Kartnig F, Pieraks M, Vladimer GI, Heinz LX, et al. LZTR1 is a regulator of RAS ubiquitination and signaling. Science. 2018;362(6419):1171–7. Epub 2018/11/18. doi: 10.1126/science.aap8210. PubMed PMID: 30442766.

29. Anderica-Romero AC, Gonzalez-Herrera IG, Santamaria A, Pedraza-Chaverri J. Cullin 3 as a novel target in diverse pathologies. Redox biology. 2013;1:366–72. Epub 2013/09/12. doi: 10.1016/j.redox.2013.07.003. PubMed PMID: 24024173; PubMed Central PMCID: PMCPMC3757711.

30. Pfeiffenberger C, Allada R. Cul3 and the BTB adaptor insomniac are key regulators of sleep homeostasis and a dopamine arousal pathway in Drosophila. PLoS Genet. 2012;8(10):e1003003. doi: 10.1371/journal.pgen.1003003. PubMed PMID: 23055946; PubMed Central PMCID: PMCPMC3464197.

31. Penev PD. Update on energy homeostasis and insufficient sleep. J Clin Endocrinol Metab. 2012;97(6):1792–801. Epub 2012/03/24. doi: 10.1210/jc.2012-1067. PubMed PMID: 22442266; PubMed Central PMCID: PMCPMC3387421.

32. Benington JH, Heller HC. Restoration of brain energy metabolism as the function of sleep. Prog Neurobiol. 1995;45(4):347-60. PubMed PMID: 7624482.

33. Petit JM, Burlet-Godinot S, Magistretti PJ, Allaman I. Glycogen metabolism and the homeostatic regulation of sleep. Metab Brain Dis. 2015;30(1):263–79. doi: 10.1007/s11011-014-9629-x. PubMed PMID: 25399336; PubMed Central PMCID: PMCPMC4544655.

34. Voll SL, Boot E, Butcher NJ, Cooper S, Heung T, Chow EW, et al. Obesity in adults with 22q11.2DS. Genet Med. 2017;19(2):204–8. Epub 2016/08/19. doi: 10.1038/gim.2016.98. PubMed PMID: 27537705; PubMed Central PMCID: PMCPMC5292049.

35. Knutson KL, Van Cauter E. Associations between sleep loss and increased risk of obesity and diabetes. Ann N Y Acad Sci. 2008;1129:287–304. Epub 2008/07/02. doi: 10.1196/annals.1417.033. PubMed PMID: 18591489; PubMed Central PMCID: PMCPMC4394987.

36. Gottesmann C. GABA mechanisms and sleep. Neuroscience. 2002;111(2):231–9. Epub 2002/05/02. PubMed PMID: 11983310.

37. Agosto J, Choi JC, Parisky KM, Stilwell G, Rosbash M, Griffith LC. Modulation of GABAA receptor desensitization uncouples sleep onset and maintenance in Drosophila. Nat Neurosci. 2008;11(3):354–9. Epub 2008/01/29. doi: 10.1038/nn2046. PubMed PMID: 18223647; PubMed Central PMCID: PMCPMC2655319.

38. Parisky KM, Agosto J, Pulver SR, Shang Y, Kuklin E, Hodge JJ, et al. PDF cells are a GABA-responsive wake-promoting component of the Drosophila sleep circuit. Neuron. 2008;60(4):672–82. doi: 10.1016/j.neuron.2008.10.042. PubMed PMID: 19038223; PubMed Central PMCID: PMCPMC2734413.

39. Chung BY, Kilman VL, Keath JR, Pitman JL, Allada R. The GABA(A) receptor RDL acts in peptidergic PDF neurons to promote sleep in Drosophila. Curr Biol. 2009;19(5):386–90. doi: 10.1016/j.cub.2009.01.040. PubMed PMID: 19230663; PubMed Central PMCID: PMCPMC3209479.

40. Potdar S, Sheeba V. Lessons from sleeping flies: insights from Drosophila melanogaster on the neuronal circuitry and importance of sleep. Journal of neurogenetics. 2013;27(1-2):23–42. doi: 10.3109/01677063.2013.791692. PubMed PMID: 23701413.

41. Gmeiner F, Kolodziejczyk A, Yoshii T, Rieger D, Nassel DR, Helfrich-Forster C. GABA(B) receptors play an essential role in maintaining sleep during the second half of the night in Drosophila melanogaster. J Exp Biol. 2013;216(Pt 20):3837–43. Epub 2013/09/27. doi: 10.1242/jeb.085563. PubMed PMID: 24068350.

42. Sheeba V, Fogle KJ, Kaneko M, Rashid S, Chou YT, Sharma VK, et al. Large ventral lateral neurons modulate arousal and sleep in Drosophila. Current biology: CB. 2008;18(20):1537–45. Epub 2008/09/06. doi: 10.1016/j.cub.2008.08.033. PubMed PMID: 18771923; PubMed Central PMCID: PMCPMC2597195.

43. Liu X, Krause WC, Davis RL. GABAA receptor RDL inhibits Drosophila olfactory associative learning. Neuron. 2007;56(6):1090–102. Epub 2007/12/21. doi: 10.1016/j.neuron.2007.10.036. PubMed PMID: 18093529; PubMed Central PMCID: PMCPMC2709803.

44. Haynes PR, Christmann BL, Griffith LC. A single pair of neurons links sleep to memory consolidation in Drosophila melanogaster. Elife. 2015;4. doi: 10.7554/eLife.03868. PubMed PMID: 25564731; PubMed Central PMCID: PMCPMC4305081.

45. Fanous AH, Kendler KS. Genetic heterogeneity, modifier genes, and quantitative phenotypes in psychiatric illness: searching for a framework. Mol Psychiatry. 2005;10(1):6–13. Epub 2004/12/25. doi: 10.1038/sj.mp.4001571. PubMed PMID: 15618952.

46. Regier DA, Narrow WE, Rae DS, Manderscheid RW, Locke BZ, Goodwin FK. The de facto US mental and addictive disorders service system. Epidemiologic catchment area prospective 1-year prevalence rates of disorders and services. Arch Gen Psychiatry. 1993;50(2):85–94. Epub 1993/02/01. doi: 10.1001/archpsyc.1993.01820140007001. PubMed PMID: 8427558.

47. Bassett AS, Marshall CR, Lionel AC, Chow EW, Scherer SW. Copy number variations and risk for schizophrenia in 22q11.2DS. Hum Mol Genet. 2008;17(24):4045–53. Epub 2008/09/23. doi: 10.1093/hmg/ddn307. PubMed PMID: 18806272; PubMed Central PMCID: PMCPMC2638574.

48. Clelland CL, Read LL, Baraldi AN, Bart CP, Pappas CA, Panek LJ, et al. Evidence for association of hyperprolinemia with schizophrenia and a measure of clinical outcome. Schizophr Res. 2011;131(1-3):139–45. Epub 2011/06/08. doi: 10.1016/j.schres.2011.05.006. PubMed PMID: 21645996; PubMed Central PMCID: PMCPMC3161723.

49. Gong L, Liu M, Jen J, Yeh ET. GNB1L, a gene deleted in the critical region for DiGeorge syndrome on 22q11, encodes a G-protein beta-subunit-like polypeptide. Biochimica et biophysica acta. 2000;1494(1-2):185–8. Epub 2000/11/10. doi: 10.1016/s0167-4781(00)00189-5. PubMed PMID: 11072084.

50. Chen YZ, Matsushita M, Girirajan S, Lisowski M, Sun E, Sul Y, et al. Evidence for involvement of GNB1L in autism. Am J Med Genet B Neuropsychiatr Genet. 2012;159B(1):61–71. Epub 2011/11/19. doi: 10.1002/ajmg.b.32002. PubMed PMID: 22095694; PubMed Central PMCID: PMCPMC3270696.

51. Ishiguro H, Koga M, Horiuchi Y, Noguchi E, Morikawa M, Suzuki Y, et al. Supportive evidence for reduced expression of GNB1L in schizophrenia. Schizophr Bull. 2010;36(4):756–65. Epub 2008/11/18. doi: 10.1093/schbul/sbn160. PubMed PMID: 19011233; PubMed Central PMCID: PMCPMC2894596.

52. Williams NM, Glaser B, Norton N, Williams H, Pierce T, Moskvina V, et al. Strong evidence that GNB1L is associated with schizophrenia. Hum Mol Genet. 2008;17(4):555–66. Epub 2007/11/16. doi: 10.1093/hmg/ddm330. PubMed PMID: 18003636.

53. Zimmerman JE, Mackiewicz M, Galante RJ, Zhang L, Cater J, Zoh C, et al. Glycogen in the brain of Drosophila melanogaster: diurnal rhythm and the effect of rest deprivation. J Neurochem. 2004;88(1):32-40. PubMed PMID: 14675147.

54. Williams JA, Su HS, Bernards A, Field J, Sehgal A. A circadian output in Drosophila mediated by neurofibromatosis-1 and Ras/MAPK. Science. 2001;293(5538):2251–6. Epub 2001/09/22. doi: 10.1126/science.1063097293/5538/2251 [pii]. PubMed PMID: 11567138.

55. Piotrowski A, Xie J, Liu YF, Poplawski AB, Gomes AR, Madanecki P, et al. Germline loss-of-function mutations in LZTR1 predispose to an inherited disorder of multiple schwannomas. Nat Genet. 2014;46(2):182–7. Epub 2013/12/24. doi: 10.1038/ng.2855. PubMed PMID: 24362817; PubMed Central PMCID: PMCPMC4352302.

56. Benes FM, Berretta S. GABAergic interneurons: implications for understanding schizophrenia and bipolar disorder. Neuropsychopharmacology. 2001;25(1):1–27. Epub 2001/05/30. doi: 10.1016/S0893-133X(01)00225-1. PubMed PMID: 11377916.

57. de Jonge JC, Vinkers CH, Hulshoff Pol HE, Marsman A. GABAergic Mechanisms in Schizophrenia: Linking Postmortem and In Vivo Studies. Front Psychiatry. 2017;8:118. Epub 2017/08/30. doi: 10.3389/fpsyt.2017.00118. PubMed PMID: 28848455; PubMed Central PMCID: PMCPMC5554536.

58. Davis RL. Mushroom bodies and Drosophila learning. Neuron. 1993;11(1):1–14. Epub 1993/07/01. PubMed PMID: 8338661.

59. Joiner WJ, Crocker A, White BH, Sehgal A. Sleep in Drosophila is regulated by adult mushroom bodies. Nature. 2006;441(7094):757–60. Epub 2006/06/09. doi: 10.1038/nature04811. PubMed PMID: 16760980.

60. Pitman JL, McGill JJ, Keegan KP, Allada R. A dynamic role for the mushroom bodies in promoting sleep in Drosophila. Nature. 2006;441(7094):753–6. Epub 2006/06/09. doi: 10.1038/nature04739. PubMed PMID: 16760979.

61. Cirelli C. The genetic and molecular regulation of sleep: from fruit flies to humans. Nature reviews Neuroscience. 2009;10(8):549–60. Epub 2009/07/21. doi: 10.1038/nrn2683. PubMed PMID: 19617891; PubMed Central PMCID: PMCPMC2767184.

62. Salkoff L, Wyman R. Genetic modification of potassium channels in Drosophila Shaker mutants. Nature. 1981;293(5829):228–30. Epub 1981/09/17. PubMed PMID: 6268986.

63. Espinosa F, Marks G, Heintz N, Joho RH. Increased motor drive and sleep loss in mice lacking Kv3-type potassium channels. Genes, brain, and behavior. 2004;3(2):90–100. Epub 2004/03/10. PubMed PMID: 15005717.

64. Kikuma K, Li X, Perry S, Li Q, Goel P, Chen C, et al. Cul3 and insomniac are required for rapid ubiquitination of postsynaptic targets and retrograde homeostatic signaling. Nature communications. 2019;10(1):2998. Epub 2019/07/07. doi: 10.1038/s41467-019-10992-6. PubMed PMID: 31278365; PubMed Central PMCID: PMCPMC6611771.

65. Roth T. A physiologic basis for the evolution of pharmacotherapy for insomnia. The Journal of clinical psychiatry. 2007;68 Suppl 5:13–8. Epub 2007/08/19. PubMed PMID: 17539704.

66. Buhr A, Bianchi MT, Baur R, Courtet P, Pignay V, Boulenger JP, et al. Functional characterization of the new human GABA(A) receptor mutation beta3(R192H). Human genetics. 2002;111(2):154–60. Epub 2002/08/22. doi: 10.1007/s00439-002-0766-7. PubMed PMID: 12189488.

67. Warming S, Costantino N, Court DL, Jenkins NA, Copeland NG. Simple and highly efficient BAC recombineering using galK selection. Nucleic Acids Res. 2005;33(4):e36. doi: 10.1093/nar/gni035. PubMed PMID: 15731329; PubMed Central PMCID: PMCPMC549575.

68. Wang S, Zhao Y, Leiby M, Zhu J. A new positive/negative selection scheme for precise BAC recombineering. Mol Biotechnol. 2009;42(1):110–6. doi: 10.1007/s12033-009-9142-3. PubMed PMID: 19160076; PubMed Central PMCID: PMCPMC2669495.

69. Pfeiffer BD, Jenett A, Hammonds AS, Ngo TT, Misra S, Murphy C, et al. Tools for neuroanatomy and neurogenetics in Drosophila. Proc Natl Acad Sci U S A. 2008;105(28):9715–20. doi: 10.1073/pnas.0803697105. PubMed PMID: 18621688; PubMed Central PMCID: PMCPMC2447866.

70. Pfeiffer BD, Ngo TT, Hibbard KL, Murphy C, Jenett A, Truman JW, et al. Refinement of tools for targeted gene expression in Drosophila. Genetics. 2010;186(2):735–55. doi: 10.1534/genetics.110.119917. PubMed PMID: 20697123; PubMed Central PMCID: PMC2942869.

71. Bischof J, Maeda RK, Hediger M, Karch F, Basler K. An optimized transgenesis system for Drosophila using germ-line-specific phiC31 integrases. Proc Natl Acad Sci U S A. 2007;104(9):3312–7. doi: 10.1073/pnas.0611511104. PubMed PMID: 17360644; PubMed Central PMCID: PMCPMC1805588.

72. Gilestro GF, Cirelli C. pySolo: a complete suite for sleep analysis in Drosophila. Bioinformatics (Oxford, England). 2009;25(11):1466–7. Epub 2009/04/17. doi: 10.1093/bioinformatics/btp237. PubMed PMID: 19369499; PubMed Central PMCID: PMCPMC2732309.

73. Klarsfeld A, Leloup JC, Rouyer F. Circadian rhythms of locomotor activity in Drosophila. Behav Processes. 2003;64(2):161–75. Epub 2003/10/15. PubMed PMID: 14556950.

